# Biomolecular condensates provide a unique environment for redox-mediated protein crosslinking

**DOI:** 10.64898/2026.04.14.718453

**Authors:** Huan Wang, Bruna Favetta, Jinying Wang, Christian Hoffmann, Elton Maloku, Yuzhou Xia, Jean Baum, Dragomir Milovanovic, Benjamin S. Schuster, Zheng Shi

## Abstract

Biomolecular condensates, often formed through liquid-liquid phase separation, are dynamic cellular compartments. Here, we demonstrate that a wide range of fluorescently tagged proteins undergo inadvertent, condensate-mediated crosslinking, resulting in rapid solidification of condensates under common fluorescence imaging conditions. The process is driven by excitation-induced, short-lived reactive oxygen species (ROS), whose otherwise limited crosslinking potential becomes uniquely enabled in the dense phase. In live cells, excitation-induced ROS potently trigger stress granule formation, while the ROS-driven solidification of condensates is modulated by compartment-dependent antioxidant buffering. Our findings demonstrate that condensates create a distinct environment that enables ROS chemistry unlikely to occur in the bulk cytosol. Furthermore, the cellular redox level can be a general regulator of condensate rheology. Beyond biological insights, our findings underscore the need for scrutiny when examining fluorophore-labeled condensates.

## Introduction

Biomolecular condensates formed by phase separation have emerged as a fundamental mechanism for organizing cellular contents. These membraneless organelles can concentrate specific biomolecules into a dense phase while still permitting dynamic exchange of components with the surroundings ^1–3^. An exciting aspect of condensates is their potential to provide a unique biochemical environment that regulates various cell signaling processes ^4–8^ or allows drug targeting ^9–12^. However, direct experimental demonstrations of the biochemical uniqueness of condensates are only emerging.

The biological functions of condensates are also intrinsically linked to their material properties, which can range from liquid-like to solid-like states ^13–16^. Condensates can mature or age, transiting from a dynamic liquid to an arrested gel or solid state ^17–20^. Solidified condensates can be functional ^21–23^, but when dysregulated, may lead to neurodegenerative diseases or cancer ^24–28^.

The investigations of condensates heavily rely on covalently tagged fluorophores, such as enhanced green fluorescent protein (EGFP), to visualize the dynamics or cellular localization of the proteins of interest ^29–32^. Excited fluorophores can enter a long-lived triplet state that inevitably generates reactive oxygen species (ROS) such as singlet oxygen, superoxide radical anion, and hydroxyl radical, inducing phototoxic effects that confound experimental results ^33, 34^. While careful protocols have been established to minimize artifacts related to fluorescence imaging ^35, 36^, potential unique challenges posed by the highly viscous, crowded biomolecular environment of a condensate have been largely overlooked.

Cellular ROS are mainly generated via the mitochondrial electron transport chain, NADPH oxidation, and peroxisomal redox metabolism ^37–39^. While excessive ROS can lead to damage or aggregation of biomolecules in ways linked to many aging-related diseases, physiological levels of ROS play critical roles in redox signaling and metabolism ^37, 38, 40^. However, the interplay between ROS and biomolecular condensates remains a major gap in our understanding of condensate pathology.

Here, we report that ROS generated by the excitation of fluorophores (e.g., EGFP) can induce extensive crosslinking across a wide range of protein condensates, resulting in a rapid, over 1000-fold increase in condensate viscosity. This phenomenon is associated with minimal photobleaching and requires the production of ROS inside the dense phase of proteins: light-induced ROS do not cause measurable crosslinking in a non-phase-separating protein solution, whereas condensates can protect proteins from crosslinking by externally added high concentrations of ROS. Fluorescence excitation also leads to the solidification of condensates in live cells. However, this effect can be reversed due to the presence of compartment-dependent antioxidants and redox buffers in cells. Furthermore, fluorescence excitation can induce the formation stress granules with properties that mimic those induced by chemical oxidants, suggesting a general role of redox potential in regulating condensate properties.

## Results

### Blue light excitation rapidly solidifies EGFP-tagged RGG condensates in vitro

The RGG domain of the P granule RNA helicase LAF-1 is a 168 residue intrinsically disordered region (IDR) that robustly forms stable liquid condensates^41^. Taking advantage of the label-free material quantification capability of micropipette aspiration (MPA)^16, 29, 42–44^, we first monitored the material properties of RGG-EGFP-RGG condensates (**Figure 1a**) under both brightfield (transmitted light that does not excite EGFP) and fluorescence (widefield blue excitation light around 480 nm) imaging conditions. During a 5-minute MPA measurement under brightfield (**Figure 1b**; white region), RGG-EGFP-RGG condensates exhibited a purely viscous behavior with a stable viscosity (η_0_) at 3.4 ± 0.3 Pa·s (mean ± SD). Surprisingly, the measured condensate viscosity (η) quickly increased upon switching to fluorescence imaging, raising over 10^4^ Pa·s after 5 minutes of continuous, low level (< 0.1 W/cm^2^) fluorescence excitation (**Figure 1b**; blue region). The condensate irreversibly transitioned from a liquid state (the condensate continuously flows with time) to a solid state (the condensate shows a stepwise deformation that does not change with time); i.e., it became effectively frozen after 5 minutes of blue light exposure (**Figure 1c; Figure S1a**). Although significant elastic features also arise with blue light exposure (**Figure 1c**), for simplicity, we focused on the quantification of viscosity and treated long-time response of an elastic condensate as infinitely viscous ^16^. Notably, while condensate viscosity rose steeply, its interfacial tension (γ) remained steady at around 100 μN/m (**Figure 1d**).

**Figure 1.**
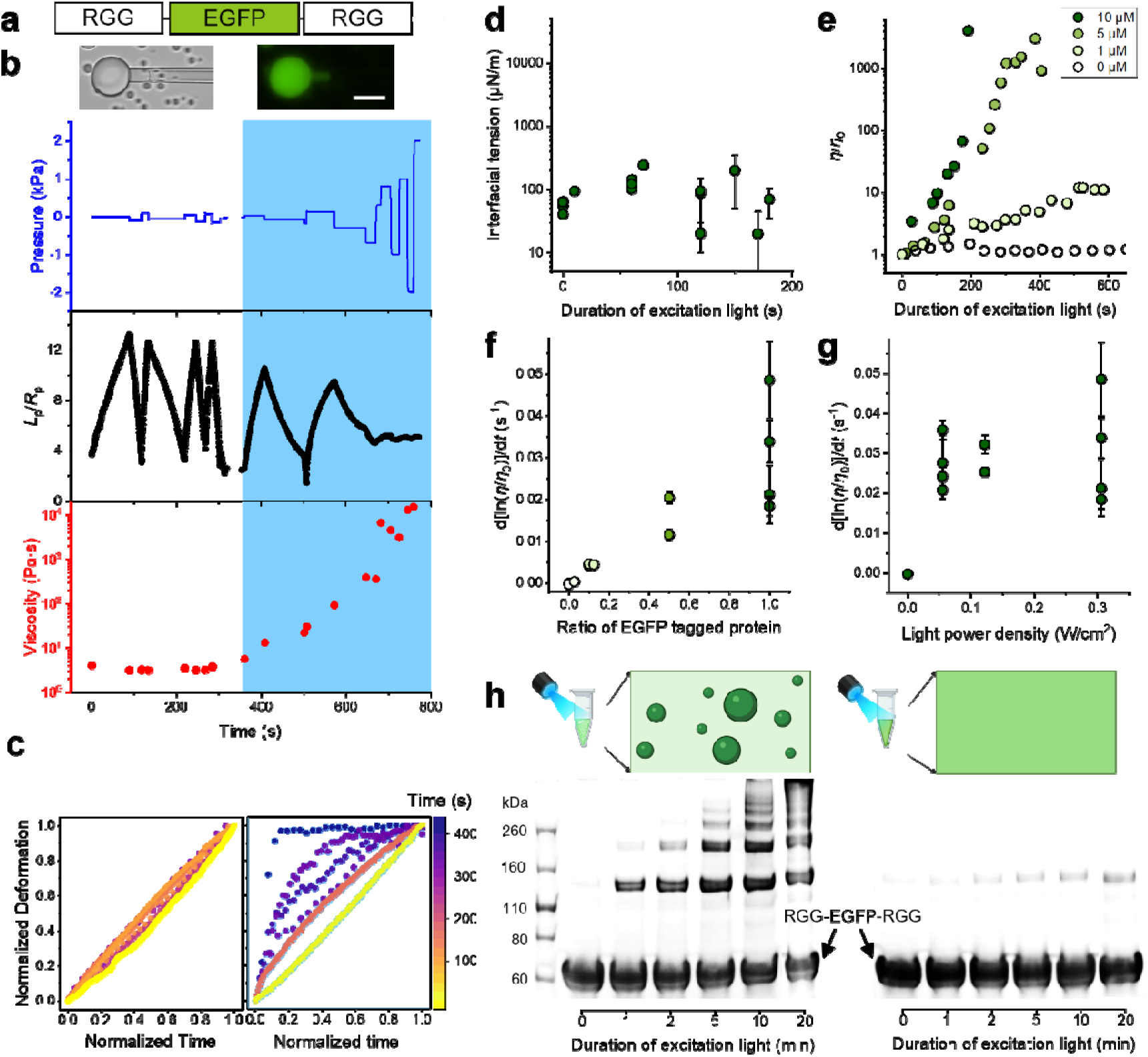
Blue light excitation leads to >1000-fold increase in the viscosity of RGG–EGFP–RGG condensates via condensate-dependent protein crosslinking. **(a)** Schematic of RGG-EGFP-RGG. All samples were 10 μM RGG-EGFP-RGG in 150 mM NaCl, 20 mM Tris, pH 7.5, 23 °C, unless specified. (**b**) Images (top) and MPA measurements (bottom) on a single condensate under brightfield (left) and fluorescence (right) imaging conditions. Blue lines: stepwise aspiration pressure. Black circles: aspiration length (*L*_p_) normalized to the micropipette radius (*R*_p_). Red circles: viscosity calculated from every pressure step. Blue excitation light was continuously on during the period marked by the blue shade. (**c**) Normalized creep curves (see Methods for definitions) for the condensate in **b** (left: brightfield; right: fluorescence). The response of a purely viscous liquid would show as a diagonal line. Departure from the line as observed in the right panel illustrates the emergence of elasticity. (**d**) Interfacial tension of the condensate vs. duration of blue excitation light. (**e**) Normalized condensate viscosity vs. duration of blue excitation light for samples containing the labeled concentrations of RGG-EGFP-RGG. All samples contain a total protein concentration of 10 μM, with the rest of the protein being RGG-RGG. The rate of condensate solidification vs. the fraction of EGFP-tagged protein in the sample (**f**) or vs. the power density of the blue excitation light (**g**). (**h**) SDS–PAGE of aliquots of the same RGG-EGFP-RGG sample under phase-separating (150 mM NaCl, 23 °C; left) or homogenous (500 mM NaCl, 23 °C; right) conditions under increasing exposure of blue light (0, 1, 2, 5, 10, 20 min). Light power density in **b**, **c**: 0.055 W/cm^2^; **d**, **e**: 0.122 W/cm^2^; **f**: 0.306 W/cm^2^; **h**: 0.133 W/cm^2^.

The blue light-induced solidification of condensates depends on the fraction of EGFP-tagged protein (RGG-EGFP-RGG), and condensates of solely unlabeled protein (RGG-RGG) did not respond to blue light (**Figure 1e**). Notably, the viscosity of condensates increased dramatically with time (**Figure 1e**), following an apparent exponential trend η = η_0_. exp(*k^RGG^_sEGFP_. t*), with the solidification rate constant *k^RGG^_sEGFP_* proportional to the fraction of RGG-EGFP-RGG in the sample (**Figure 1f**). These measurements suggest a key role of EGFP excitation in the observed solidification of condensates. The solidification rate of pure RGG-EGFP-RGG condensates depends weakly on the power density of blue light between 0.05 – 0.3 W/cm^2^ (**Figure 1g**), a range commonly used to image EGFP and induces minimal photobleaching (**Figure S1b**) ^45, 46^.

Viscosity increase of a polymer solution is often a result of molecular crosslinking, with ‘ln η’ proportional to the average size of the crosslinked polymer^47^. A fully crosslinked polymer solution can form a system-spanning network and behave as a viscoelastic solid with an infinite viscosity at long-time^48^. Indeed, from SDS-PAGE analysis of phase-separated RGG-EGFP-RGG samples (150 mM NaCl, 23 °C) under blue light exposure, we observed clear dimers and higher-order oligomers of RGG-EGFP-RGG, with their abundance increasing proportionally with the duration of light (**Figure 1h**, left). Surprisingly, blue light did not induce detectable oligomers of RGG-EGFP-RGG when protein phase separation was inhibited with either elevated salt (500 mM NaCl) or temperature (42 °C) (**Figure 1h**, right**; Figure S1c**). These results directly demonstrate that condensates provide a unique biochemical environment that allows irreversible crosslinking of RGG-EGFP-RGG. Under phase-separating conditions, proteins stay monomeric in both RGG-EGFP-RGG (i.e., labeled) samples kept in the dark (**Figure S1c**) and RGG-RGG (i.e., unlabeled) samples exposed to blue light (**Figure S1d**), demonstrating that EGFP excitation is essential for inducing condensate-specific crosslinking.

### Excitation-induced solidification is a general property of fluorophore-containing condensates

To test whether the phenomenon of excitation-induced condensate solidification is unique to RGG-EGFP-RGG, we next examined a panel of phase-separating proteins where either the EGFP or the phase-separating RGG domains were substituted with other proteins or fluorophores (Figure 2a).

**Figure 2.**
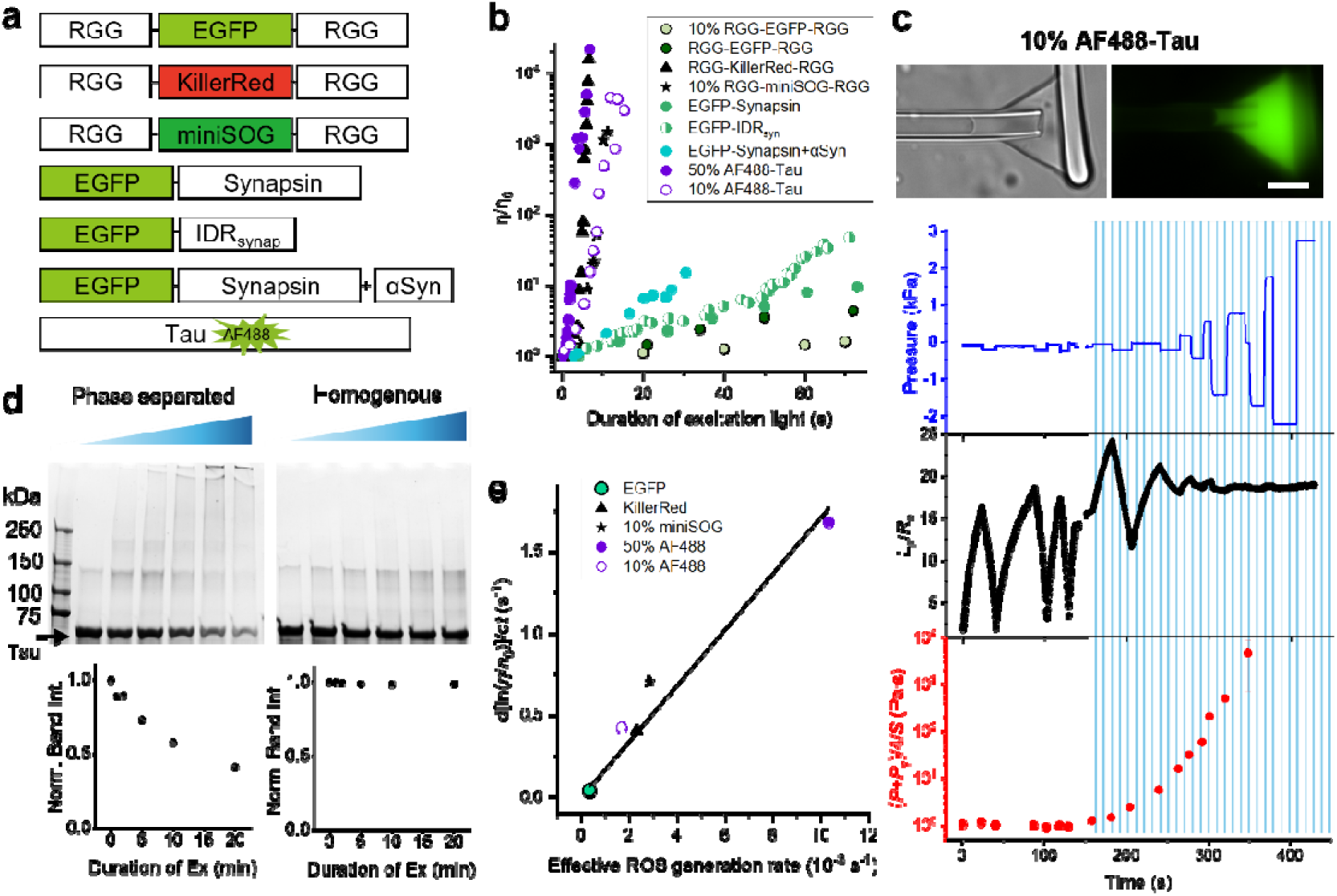
Excitation induced condensate solidification observed across a wide range of protein condensates. **(a)** Schematics of studied proteins. Fluorophores were incorporated either genetically (EGFP, KillerRed, miniSOG) or by covalent dye attachment (AF488) to phase-separating scaffolds: RGG-RGG, Synapsin (full-length or IDR), or Tau. (**b**) Excitation induced condensate viscosity increase for each protein in **a**, plotted as the relative viscosity (η/η₀, in log scale) vs. the duration of fluorescence excitation. (**c**) A representative MPA measurement on AF488-labeled Tau condensates (10% labeling) similar to that in Figure 1b, except under intermittent excitation (60 ms of blue light per second). (**d**) SDS–PAGE analysis of phase-separated (with 10% PEG) vs. homogeneous (no PEG) Tau samples (20 μM Tau, 10% labeling) exposed to increasing light doses (1, 2, 5, 10, 20 minutes blue light). The gray curve is an exponential fit (R^2^ = 0.99), with decay time (10 ± 2) min and saturation intensity 0.31 ± 0.07. (**e**) Condensate solidification rates scale linearly with the effective ROS generation rate, which is the product of molar fraction of fluorescent labeling and the bleaching rate of the observed fluorescence. Black line was a linear fit. RGG based samples were all 10 μM in 150 mM NaCl, 20 mM Tris, pH=7.5. Tau samples were 20 μM in 10% PEG-8000, 25LJmM HEPES, 150 mM NaCl, pH=7.4. Synapsin samples were 9 μM with or without 3 μM alpha-synuclein in 3% PEG-8000, 150 mM NaCl, 25 mM Tris, and 0.5 mM TCEP, pH=7.4. Synapsin IDR samples were 9 μM in 10% PEG-8000, 150 mM NaCl, 25 mM Tris-HCl, and 0.5 mM TCEP, pH=7.4. KillerRed was excited under green light with a power density of 0.314 W/cm^2^, all other proteins were excited under blue light with a power density of 0.306 W/cm^2^ (**b**, **c**, **e**) or 0.133 W/cm^2^ (**d**)

Considering the potential involvement of excitation-induced ROS, we replaced EGFP with KillerRed or miniSOG—red and green fluorescent proteins, respectively—both known for their strong ROS generation capacity and phototoxicity ^49, 50^. Both RGG-KillerRed-RGG and RGG-miniSOG-RGG condensates rapidly solidified under fluorescence excitation (Figure 2b**, Figure S2**). When excited by green light, the solidification rate of RGG-KillerRed-RGG condensates (*k^RGG^_sKillerRed_*) was ∼20 times that of RGG-EGFP-RGG (*k^RGG^_sEGFP_*), under equal power density of blue light). Under blue light excitation, condensates of solely RGG-miniSOG-RGG solidified too rapidly for accurate viscosity tracking via MPA. Reducing the labeling fraction to 10%, we found *k^RGG^_sminiSOG_* ∼100-fold greater than *k^RGG^_sEGFP_* under the same condition. The comparisons between EGFP with KillerRed and with miniSOG directly suggest that excitation-induced ROS, rather than the excitation lights themselves, being the direct cause of condensate solidification.

We next swapped the RGG domains with Synapsin, a major presynaptic protein whose phase separation is driven by its C-terminal IDR ^51, 52^. EGFP labeled condensates of both full length Synapsin (*k^Synapsin^_sEGFP_*) and its IDR (k^IDR^*_sEGFP_*) showed significant viscosity increase upon exposure to blue light, with solidification rates ∼2-fold of k^RGG^*_sEGFP_* under similar sample and imaging conditions (Figure 2b**, Figure S2**). These results suggest that excitation induced condensate solidification is a general phenomenon, governed primarily by the ROS-generating capacity of the fluorophore and less sensitive to the phase-separating protein scaffolds (**Table S1**)

Studies of condensate solidification have largely focused on proteins linked to aging-related diseases^1, 53, 54^. Therefore, we analyzed two proteins closely associated with neurodegeneration: Tau and alpha-synuclein (αSyn) ^55, 56^. We studied Tau condensates labeled with Alexa Fluor™ 488 (AF488) to also assess whether small-molecule dyes exhibit properties distinct from those of large fluorescent proteins. Due to αSyn’s weak propensity to form condensates on its own ^55^, we studied αSyn as a client that partitions into EGFP-Synapsin condensates. The presence of low levels of unlabeled αSyn (3 µM) increased the solidification rate of Synapsin condensates by 3-fold (Figure 2b**, Figure S2**), consistent with αSyn’s ability to increase the viscosity and speed up the maturation of Synapsin condensates in vitro and in live cells^16^. Condensates containing AF488-labeled Tau rapidly solidified under blue light. At comparable protein labeling fraction, k^Tau^_sAF488_ was ∼100-fold greater than k^RGG^_sEGFP_ (Figure 2b). The accelerated solidification of AF488-Tau condensates is consistent with the high phototoxicity of small molecule dyes ^57^, and may be further enhanced by the intrinsic fibrillation propensity of Tau.

Figure 2c shows a representative MPA experiment for Tau condensates with 10% labeling. Like RGG-EGFP-RGG condensates (Figure 1), the viscosity of Tau condensates remained stable under brightfield but increased rapidly upon exposure to blue light. To allow accurate MPA-based viscosity tracking and to better mimic typical condensate microscopy experiments, we applied blue light intermittently only during imaging (60 ms of blue light per second). The results suggest that minimal, non-continuous, excitation light is sufficient to trigger the solidification of condensates.

Consistent with the condensate-dependent crosslinking of RGG-EGFP-RGG (Figure 1h), SDS-PAGE analysis confirmed the formation of higher-order oligomers in phase-separated (10% PEG), but not homogeneous (no PEG) AF488-Tau samples (Figure 2d). Notably, even though the sample only contained 10% of labeled Tau, 60 % of all Tau were oligomerized after 20 min of blue light exposure, suggesting that crosslinking can happen to all proteins in the condensates, not just the ones labeled with the fluorophore. Since the crosslinking of Tau only happens in the dense phase, the amount of Tau monomers after extensive blue light directly reports Tau concentration in the dilute phase (i.e., the saturation concentration for phase transition: *c*_sat_). For the 10 μM sample shown in Figure 2d, we found *c*_sat_ = 3.1 ± 0.7 μM, consistent with direct measurements of the phase diagram of Tau^58^.

The generation of ROS from the excitation of a fluorophore is directly correlated to the fluorophore’s photostability ^59^. Therefore, we compiled the solidification rates (*k*_s_) of all studied condensates according to an effective ROS generation rate in each sample based on its photobleaching rate (*k*_pb_) and labeling fraction (¢; Figure 2e). A linear relation was observed between *k*_s_ and ‘ ¢ *k*_pb_’ (slope = 171 ± 12; R^2^ = 0.98), further supporting that the observed solidification of condensates is universal to fluorescently labeled phase-separating proteins and closely associated with excitation-induced ROS generation. Importantly, Figure 2e provides a practical framework for estimating the impact of fluorescence imaging on condensate viscosity:, a ∼4000-fold increase in condensate viscosity is expected for every 5% decay in the fluorescence intensity (*I*) of a fully labeled condensate (see Supplementary Discussion). Notably, this relation also enables the estimation of the fluorescence labeling fraction () of condensates with unknown composition, a capacity particularly valuable for investigating condensates in cells and in vivo.

### The solidification of condensates is driven by fluorophore-generated ROS within the condensate

To delineate the chemical mechanism responsible for light-induced crosslinking, we focused on examining condensates of RGG–EGFP–RGG due to its simplicity in protein sequence (no cysteine in the LAF-1 RGG domain; **Table S1**), buffer condition (no crowder needed for condensate formation), and capacity to form large and stable condensates (condensates maintain liquidity for days without fluorescence excitation) ^60^. We found that fluorescence recovery after photobleaching (FRAP), a widely used technique to estimate molecular mobility in condensates ^61^, can reliably track excitation-induced increase in the viscosity of RGG condensates up to ∼ 500 Pa·s (**Figure S3**). Therefore, next we used FRAP as a broadly accessible tool to demonstrate the relevance of our previous findings, and to screen factors that modulate light-induced solidification of RGG condensates.

Short exposure to blue light (equivalent to < 5 s of continuous, 0.122 W/cm^2^, 488 nm light) during one round of FRAP measurement has minimal effect on the measured mobility of RGG–EGFP–RGG, with the bleached region reaching ∼90% recovery after 40 s (Figure 3a). However, extended exposure (> 1 min) to blue light before the FRAP experiment leads to consistently slower recovery of the bleached region, with recovery at 40 s reduced to ∼10% after 5 min of blue light exposure (Figure 3b), consistent with MPA-based viscosity measurements (Figure 1). In contrast, no significant decrease in the recovery fraction was observed when exposing the condensate to 5 min of green light (0.182 W/cm^2^, 561 nm), directly confirming the essential role of fluorescence excitation in immobilizing proteins in the condensate (Figure 3c).

**Figure 3.**
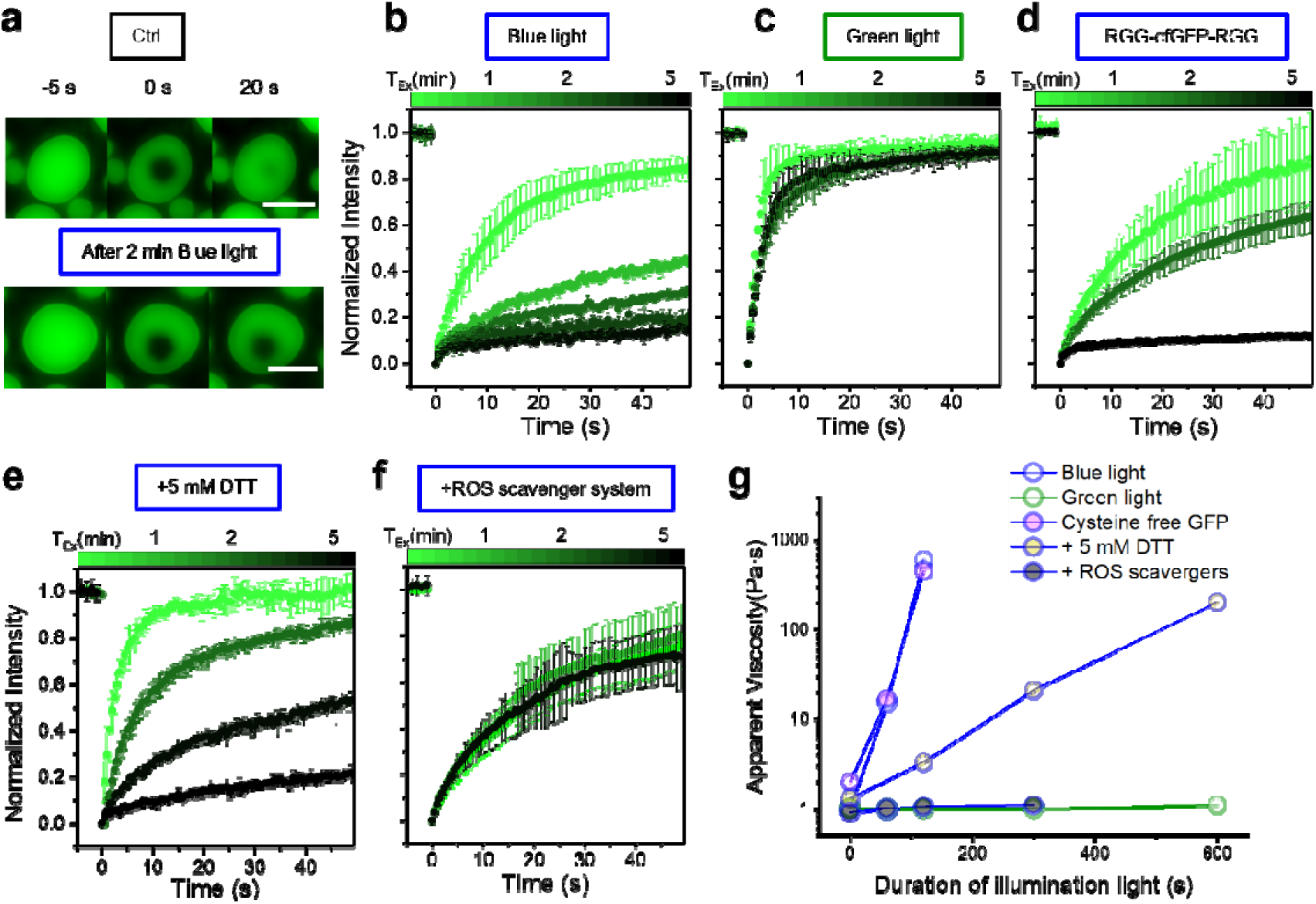
The solidification of condensates is caused by ROS generated during fluorescence excitation. **(a)** Representative FRAP experiments of an RGG-EGFP-RGG condensate before (top) and after 2 minutes of continuous blue light exposure (bottom). Scale bars, 5 µm. (**b**) FRAP experiments on RGG-EGFP-RGG condensates exposed to increasing amount of blue light (488 nm). Line color indicates T_EX_, the duration of excitation light exposure, with darker colors corresponding to longer duration. (**c-f**) Same as **(b)** except blue light was switched to green light (561 nm) in (**c**); (**d**) added 5 mM DTT in the buffer; (**e**) EGFP substituted to a cysteine-free EGFP mutant (cfGFP); (**f**) added an ROS scavenging system. (**g**) Apparent condensate viscosity calculated from FRAP vs. illumination time for conditions in (**b**) to (**f**). Power density of blue light in figure **a**, **b**, **d**, **e**, **f, g**: 0.122 W/cm^2^. Power density of green light in figure **c** and **g**: 0.182 W/cm^2^.

RGG domains do not contain any cysteine, the residue with the lowest reduction potential and most highly susceptible to crosslinking through the formation of disulfide bond ^62^. Therefore, replacing EGFP with cysteine-free GFP (cfGFP; C48S and C70M) creates a phase-separating protein scaffold that enables the direct assessment of molecular contributions to condensate solidification beyond cysteine crosslinking. RGG-cfGFP-RGG condensates showed phase behavior and blue light response indistinguishable from its EGFP counterparts (Figure 3d). Furthermore, we use dithiothreitol (DTT) to create a reducing environment and prevent the formation of disulfide bonds ^63^. RGG-EGFP-RGG condensates still exhibit excitation-induced solidification in the presence of 5 mM DTT, but at a prominently slower rate (Figure 3e). These results indicate that excitation-induced crosslinking, while dependent on the redox potential of the environment, is not restricted to cysteine residues.

RGG domains contain a large fraction of tyrosine (11 out of 168 amino acids), the residue with the second lowest reduction potential ^64^. The formation of dityrosine would lead to strong light absorbance at 315 nm ^65^. Monitoring the UV-vis spectrum of RGG-EGFP-RGG condensates exposed to blue light, we observed a time-dependent increase of 315 nm absorbance, accompanied by a decrease of absorbance at 488 nm (**Figure S4**). This result suggests the formation of tyrosine–tyrosine crosslinking in response to excitation-induced photobleaching of EGFP ^66^.

Finally, we introduced a ROS-scavenger system (7.5 U/mL pyranose oxidase, 1 kU/mL catalase, and 50 mM glucose) ^35^ to completely remove ROS in the sample. The ROS-scavenger system, which likely partitions into the condensates, abolished excitation-induced solidification of RGG-EGFP-RGG condensates (Figure 3f). Collectively, these results (Figure 3g) demonstrate that fluorophore excitation-generated ROS drive protein crosslinking within condensates beyond simple disulfide bond formation (e.g., dityrosine).

### Condensates can protect proteins from getting crosslinked by external ROS

Excitation-induced ROS are generated throughout the fluorophore-tagged, phase-separating protein sample. To examine how the spatial distribution of ROS influences the crosslinking of phase-separating proteins, we introduced hydroxyl radicals (**·**OH, a highly reactive ROS), via the Fenton reaction (10 mM H_2_O_2_ + 1 mM FeCl_2_)^67^ into 10 μM RGG-EGFP-RGG samples in either a homogeneous state (42 °C) or a phase-separated state (23 °C). In both cases, FeCl_2_ was added to protein samples premixed with H_2_O_2_ (Figure 4a).

**Figure 4.**
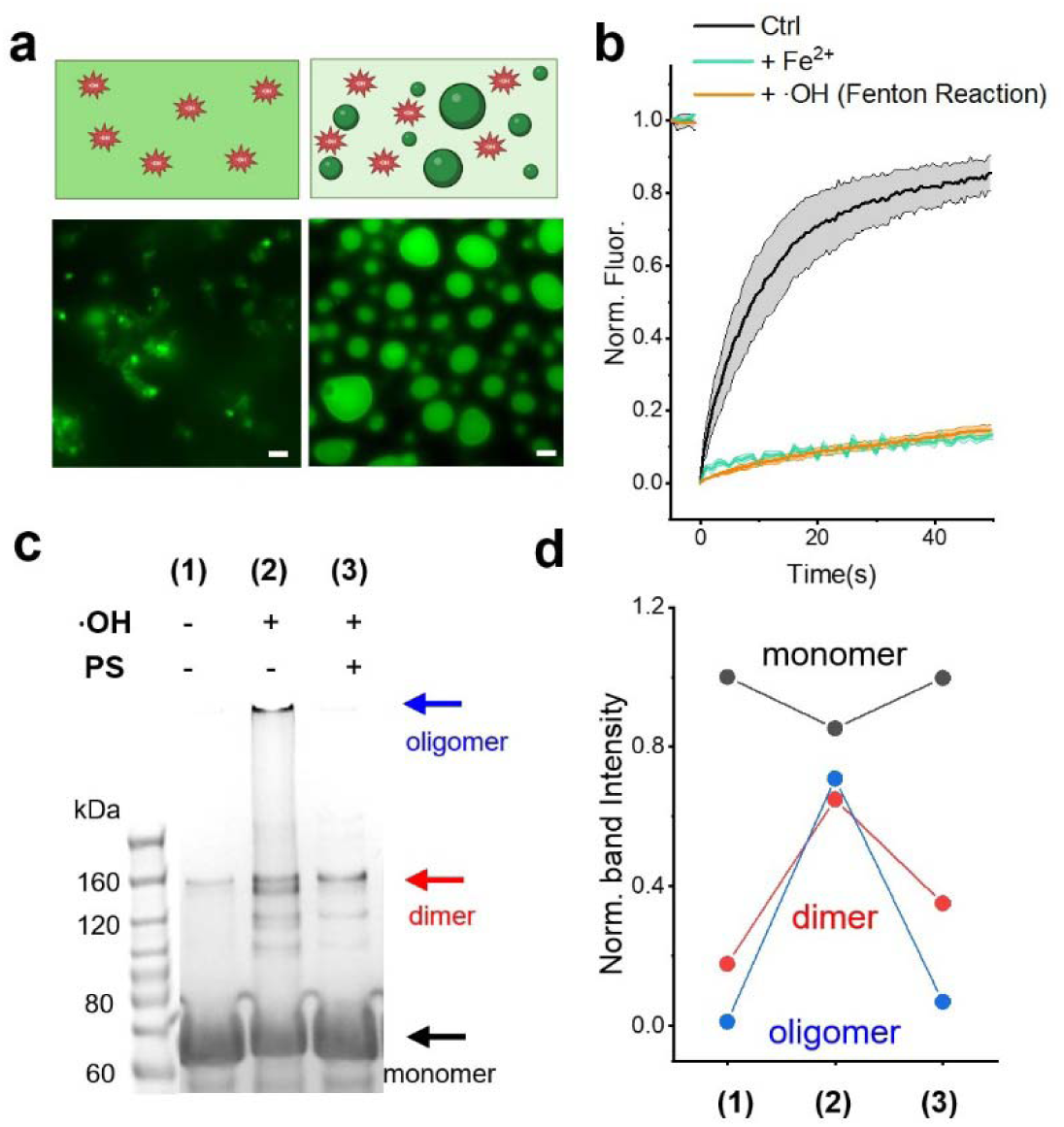
Response of RGG-EGFP-RGG to externally added hydroxyl radicals. (**a**) Schematics (top) and fluorescence images (bottom) after adding hydroxyl radicals (from Fenton reaction) to 10 μM RGG-EGFP-RGG in a homogeneous state (42 °C, left) or a phase-separated state (23 °C, right). Scale bars, 5 µm. (**b**) FRAP experiments on condensates of 10 μM RGG-EGFP-RGG at 23 °C (Ctrl). ‘+ Fe^2+^’: add 1 mM FeCl_2_ to the control sample. ‘+ **·**OH’: add 1 mM FeCl_2_ to the control sample premixed with 10 mM H_2_O_2_. SDS-PAGE image (**c**) and quantification (**d**) of RGG-EGFP-RGG samples with or without **·**OH under homogeneous (-PS) or phase-separated (+PS) conditions.

RGG-EGFP-RGG contains a C-terminal His-tag (**Table S1**) capable of coordinating with Fe^2+^ ^68^. In line with this property, the addition of FeCl_2_ significantly reduced the mobility of proteins in the dense phase (Figure 4b). We hypothesize that the coordinated trapping of Fe^2+^ by His-Tag and the highly viscous environment inside the condensate (∼3000-fold higher than the viscosity of the dilute phase) would effectively inhibit the Fenton reaction^69^, leaving the production of ·OH mainly in the dilute phase. With a half-life ∼1 ns ^70^, **·**OH formed in the dilute phase (characteristic diffusion length ∼5 nm in the dilute phase and ∼0.1 nm in condensates) would not penetrate the outer surface layer of the condensates. Consistent with this hypothesis, the addtion of H_2_O_2_ + FeCl_2_ (i.e., prodcution of **·**OH) did not reduce the mobility of proteins inside the condensate beyond that of FeCl_2_ itself (Figure 4b).

Through microscopy and SDS-PAGE, we found that the high concentration of **·**OH produced from the Fenton reaction caused pronounced aggregation and extensive cross-linking of the homogeneous protein sample (Figure 4a**, 4c**). However, the amount of **·**OH-induced irreversible protein aggregation was significantly reduced in the phase-separated sample (Figure 4c**, 4d; Figure S5**). This result further supports the key role of ROS in the previously observed protein crosslinking and condensate solidification. More importantly, it suggests that condensate formation can serve as a protective mechanism that mitigates protein crosslinking caused by externally added ROS.

### ROS-driven solidification of condensates is buffered in cells

We next sought to determine if the ROS-induced solidification of in vitro condensates could also occur in the complex environment of a live cell. We first focused on stress granules (SGs), a well-studied type of cytosolic condensate that directly interplays with cellular ROS levels. Cells assemble G3BP1-enriched SGs in response to oxidative, pH, osmotic stresses, or the overexpression of G3BP1 ^71^. To achieve intracellular quantification, we performed micropipette aspiration and whole-cell patch clamp (MAPAC)^16^ measurements on SGs induced by overexpressing EGFP-G3BP1 in HEK293T cells (Figure 5a). While fluorescence excitation is necessary for monitoring intracellular condensates, we limited the excitation to a minimal level (∼0.1 W/cm^2^ power density and 10-30 s total exposure) during MAPAC. Then, we used a prolonged ∼0.6 W/cm^2^ blue light to evaluate the effect of excitation-induced ROS.

**Figure 5.**
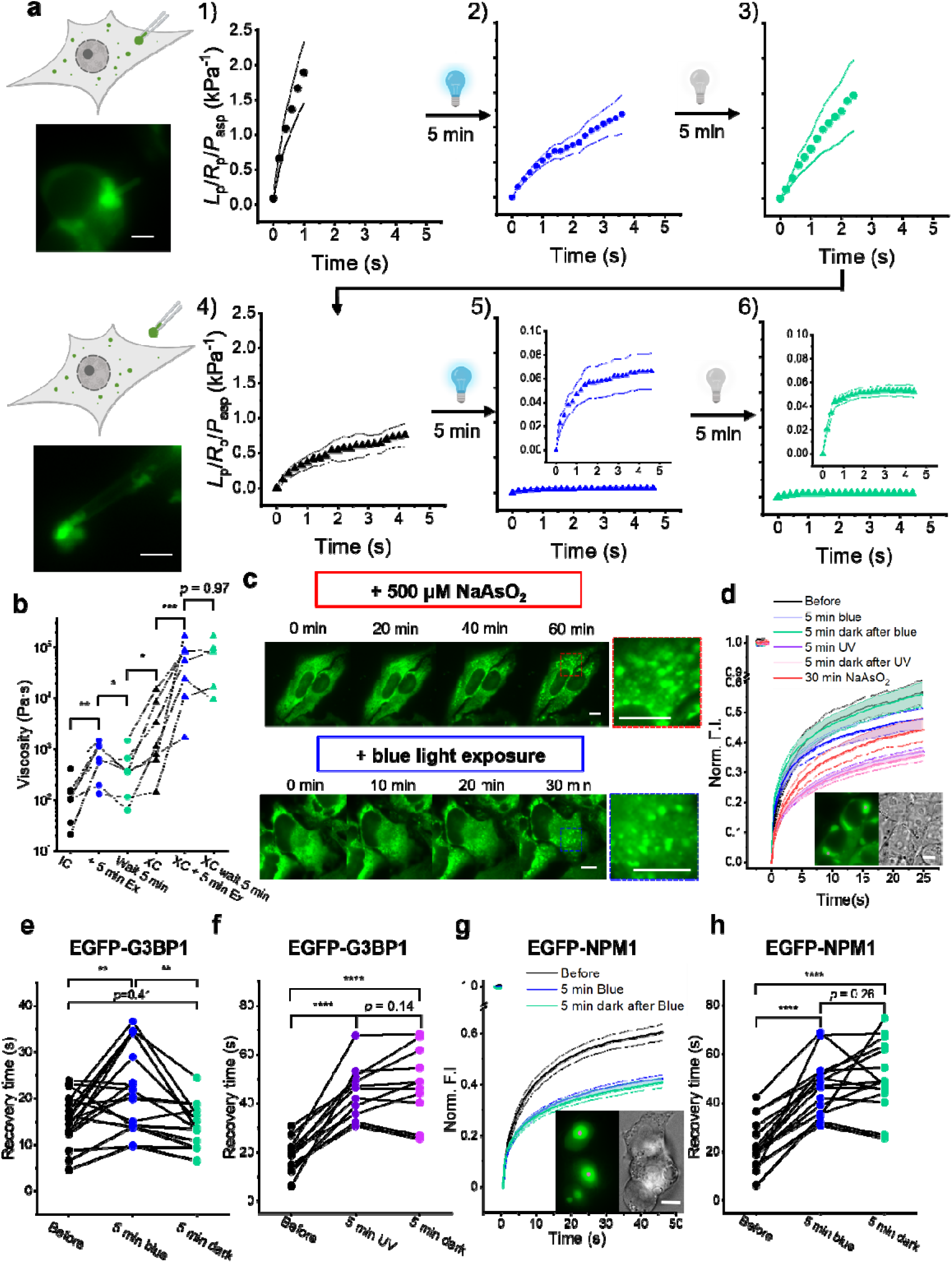
Fluorescence excitation leads to viscosity increase of cellula condensates, buffered by cytosolic antioxidants. **(a)** MAPAC measurements of SGs in HEK293T cells expressing EGFP-G3BP1. Left: schematics and fluorescence images showing an aspirated intracellular (up) and extracted (down) SG (scale bars, 2 µm). Right: normalized condensate deformation 1) before illumination (black), 2) after 5 min blue light exposure, 3) after 5 min recovery in the dark, 4) after extracting the condensate to extracellular medium, 5) after 5 min blue light exposure on the extracted condensate, 6) after 5 min recovery in the dark. Light power density during imaging: 0.122 W/cm^2^. Light power density for the 5-minute blue light exposure: 0.596 W/cm^2^. (**b**) MAPAC-quantified viscosity of stress granules from 7 cells across conditions listed in **a**. (**c**) Time-lapse fluorescence images of HeLa cells expressing EGFP-G3BP1. Top: SGs induced by 500 µM NaAsO_2_. Bottom: SGs induced by continuous exposure to 0.063 W/cm^2^ blue light. Imaging condition:100 ms exposure per min. Scale bars, 10 µm. (**d**) FRAP measurements of EGFP-G3BP1 in SGs under the following conditions: before any treatment (black); after 5 minutes of 0.974 W/cm^2^ blue light exposure (blue); after 5 minutes recovery in the dark following the blue light exposure (green); after 5 minutes 0.118 W/cm^2^ UV exposure (purple); after 5 minutes recovery in the dark following the UV exposure (pink); after 30 minutes incubation in 500 µM NaAsO_2_(red). (**e**-**f**) FRAP recovery time of EGFP-G3BP1 measured from SGs in HEK293T treated with 5 min blue light (**e,** n = 18) or UV (**f,** n = 11), *p* value reported from paired t-test. (**g**) FRAP measurements of EGFP-NPM1 expressed in HEK293T cells (n = 15). (**h**) Recovery time for conditions in **g**. *p* value reported from paired t-test. In this figure, \**p* < 0.05, \*\**p* < 0.01, \*\*\**p* < 0.001, \*\*\*\**p* < 0.0001.

As demonstrated in Figure 5a and quantified in Figure 5b, SGs flow easily into the micropipette before prolonged blue light, corresponding to a viscosity of 100 Pa·s (median; same below). This viscosity resembles that of fluid condensates in live cells^16^. After 5 minutes of blue light exposure, the viscosity of SGs increased to 600 Pa·s. The solidification of SGs was significant (*p* < 0.01), with a rate comparable to those observed on reconstituted condensates containing 10% EGFP-labeled proteins (Figures 1e**, 1f**). However, in contrast to the irreversible solidification of in vitro condensates, the viscosity of SGs reduced to 350 Pa·s after a 5-minute recovery period in the dark, suggesting the presence of antioxidants in the cytosol (e.g., glutathione, superoxide dismutase, catalase, glutathione peroxidase) that can buffer or break ROS-induced biomolecular crosslinking^72^. To test this, we extracted the measured SGs to the extracellular medium. Extracted SGs became substantially more viscous (1 kPa·s), reflecting their transition from the reducing cytosol to the more oxidative extracellular environment. Furthermore, 5 minutes of blue light induced a significant increase in the viscosity of extracted SGs (60 kPa·s; *p* < 0.001) that maintained after the 5-minute recovery period in dark (*p* = 0.97), consistent with the lack of antioxidants in the extracellular medium. The extracted SGs showed strong elastic features (i.e., no measurable flow at long time) upon blue light (**Figure 5a5-6**), reminiscent of those observed on solid cellular condensates that are associated with diseases ^16^.

The solidification of SGs suggests that excitation-induced ROS can also crosslink proteins in complex cellular condensates. Consistently, in HeLa cells that exhibit diffusive EGFP-G3BP1 signals, continuous blue light exposure was sufficient to induce the formation of microscale SGs that were morphologically indistinguishable from those induced by the commonly used chemical oxidant NaAsO_2_ (Figure 5c) ^73^. Moreover, the mobility of EGFP-G3BP1 in cellular SGs exposed to prolonged blue light was comparable to that in NaAsO_2_-induced SGs (Figure 5d).

Compared to extracted or in vitro condensates, a distinguishing feature of intracellular SGs is their capacity to reverse excitation-induced solidification (**Figure 5a1-3**). We next used FRAP as a less invasive assay to verify the resilience of intracellular SGs (Figure 5d). High levels of blue light led to a significant reduction in the mobility of EGFP-G3BP1 in SGs (Figure 5e), but not in the dilute phase (**Figure S6a**). This condensate-specific effect requires sufficient EGFP excitation within the dense phase —neither low-level excitation of condensates (**Figure S6b**) nor high-level excitation of EGFP in the dilute phase (**Figure S6c**) altered the mobility of fluorescently tagged G3BP1. These observations echo those made on reconstituted condensates (Figures 1-3) and are consistent with MAPAC measurements of condensates in cells (Figure 5a**, 5b**). More importantly, excitation-induced immobilization of EGFP-G3BP1 in SGs fully reversed following a 5-minute dark recovery period (Figure 5d**, 5e**), confirming that an intact intracellular environment permits efficient buffering of ROS-induced protein crosslinking.

We hypothesize that antioxidants in the cell played a key role in counteracting ROS-induced condensate solidification. To test this, we use UV irradiation to overwhelm the redox system in cells. We observed that high dosage of UV exposure efficiently kills cells, while low level of UV light induces SG formation, accompanied by a significant, long-lasting increase in the ROS level across the entire cytoplasm (**Figure S7**). In cells containing pre-existing SGs, a 5-minute UV exposure significantly reduced the mobility of EGFP-G3BP1 in SGs (Figure 5d**, Figure S6d**). In contrast to the transient effect induced by blue light, the UV-induced reduction in EGFP-G3BP1 mobility persisted after a 5-minute dark period (Figure 5d**, 5f**), consistent with the depletion of intracellular antioxidants. A more oxidizing cytoplasm is often associated with depolarized cell membrane potential^74^. Indeed, membrane voltage recordings in MAPAC showed that SGs tend to be more liquid in cells with a more polarizing membrane potential (**Figure S8**).

Most antioxidant enzymes in cells are in the cytosol^75^, therefore, we expect nuclear condensates to be less resilient to excitation-induced solidification. Using EGFP-NPM1 to label the nucleoli, one of the largest cellular condensates ^16, 76^, we observed a significant reduction in the mobility of NPM1 upon blue light (Figure 5g). Under the same blue light level, the immobilization of EGFP-NPM1 was more prominent than that of EGFP-G3BP1 (Figure 5h). More importantly, EGFP-NPM1 maintained low mobility after the 5-minute dark period (Figure 5g**, 5h**), consistent with the lack of antioxidants in the nucleoplasm.

Taken together, our findings demonstrate that fluorescence excitation can induce the solidification of condensates in live cells. However, the intracellular environment provides a significant buffering effect that varies according to the cellular state and compartment, illustrating how intracellular redox levels can directly regulate condensate rheology.

## Discussion

Our results reveal that biomolecular condensates provide a unique environment that regulates ROS-mediated protein crosslinking. Excitation of fluorophore-tagged condensates under common imaging conditions can lead to dramatic, >1000-fold increases in condensate viscosity, converting liquid droplets into viscoelastic or solid-like states. This crosslinking relies on the formation of a dense phase (Figure 1h) and is a general property of fluorophore-containing condensates, with solidification rates directly correlated with the fluorophore’s capacity to produce ROS (Figure 2e). Previous studies exploring the biochemical uniqueness of condensates have largely focused on the partitioning of stable molecules ^9–12^. However, for short-lived species such as ROS, the dense biomolecular environment in a condensate can lead to reactions that would not happen in a dilute condition.

We also report a dual role of condensates in regulating redox chemistry. Condensates promote protein crosslinking when the ROS source is internal (i.e., from tagged proteins within the dense phase; Figures 1-3) but can also protect proteins from externally added ROS (Figure 4). This protective effect likely arises from the high viscosity within the condensate, which can limit the diffusion of external ROS and deplete free oxygen inside condensates ^77^. Depending on the cellular context, condensates may be used to promote specific protein crosslinking or to protect biomolecules from oxidative damage.

Our findings have profound methodological implications. The “aging” or “maturation” of condensates, often characterized by a time-dependent decrease in FRAP recovery or an increase in viscosity, is widely studied due to its potential connection to diseases ^2, 18, 28^. Our results caution that, in many in vitro systems, the apparent maturation of condensates is confounded by strong excitation-induced artifacts. The very act of measuring condensate material property based on fluorescence excitation could be the main driver of condensate maturation, highlighting a critical need for researchers to employ label-free techniques^78^. When fluorescent imaging is inevitable, one should select fluorophores with minimal ROS-generation capacity, minimize the fraction of labeling, and reduce the power and frequence of excitation light (see Supplementary Discussion), in addition to considering steric effects of fluorescent tagging^79^.

Interestingly, our study also indicates that the special biochemical environment inside a condensate could be utilized for local compositional analysis both in vitro and in cells. By triggering the local generation of ROS and subsequent crosslinking within the condensate, researchers could map the proteome of specific condensates with high spatial resolution. This conceptually mirrors existing proximity labeling methods ^80, 81^, but obviates the use of potentially toxic strong ROS generators.

Our findings also have broad implications in understanding the role of cellular condensates in regulating oxidative stress and DNA damage. We show that fluorescence excitation can induce the solidification of EGFP-tagged condensates in live cells. In contrast to reconstituted condensates, the solidification of condensates in healthy cells can be strongly buffered and even reversed (Figure 5). This demonstrates the power of the cytoplasmic redox buffering system, which can quench excess ROS and even reverse protein crosslinking, resulting in a homeostasis of condensate fluidity despite oxidative perturbation. However, when this system is overwhelmed, such as cells exposed to extensive UV irradiation, condensates can be irreversibly solidified. These observations establish a direct link between cellular health, redox homeostasis, and condensate fluidity, suggesting that the pathological solidification observed in disease may represent a failure of these protective buffering mechanisms.

Several limitations remain in this work. While we tested a range of fluorophores and phase-separating proteins, the susceptibilities of many other biomolecular condensates and commonly used fluorophores remain unexplored. We focused on crosslinking between proteins; however, an extensive mapping of crosslinking sites and the potential involvement of protein folding ^82^ or other biomolecules^6, 8^ are not considered. Notably, a recent report indicates that UV-induced crosslinking alters the material state of DNA/RNA-based condensates ^83^. Lastly, we inferred the role of cellular antioxidants, but real-time monitoring of the absolute ROS level within cellular condensates remains challenging. Future studies utilizing high specificity, ratiometric ROS sensors could provide key insights in the relation between condensate biology and intracellular redox signaling.

## Acknowledgements

We thank funding supports by the National Institutes of Health (NIH) grant R35GM147027 and National Science Foundation (NSF) grant MCB-2440729 to Z.S.; NIH grant R35GM142903 and NSF grant DMR-2238914 to B.S.S.; NIH grant R35GM136431 to J.B.; German Center for Neurodegenerative Diseases (DZNE) start-up funds, European Research Council Grant MemLessInterface (101078172) and Horizons Europe MSCA Doctoral Network (101167843 - BICEPS) to D.M..

## Author Contributions

Z.S. and H.W. conceived the project and designed the experiments. H.W. performed the core experiments and analyzed the data. B.F., J.W., and C. H. purified the proteins in this study. E.M., Y.X. helped the experiments in Fig 5c. H.W. wrote the initial manuscript. H.W., B.S.S., and Z.S. discussed about the results and revised the manuscript. All authors commented on the manuscript.

## Methods

### Protein expression and purification

The expression and purification of RGG based proteins, EGFP-Synapsin 1, alpha-Synuclein are the same as previously described^16, 42^.

The expression and purification of Tau: The genes of WT 2N4R human tau (441 residues, without His-Tag) were subcloned into a pET28 plasmid to express intact tau protein in Escherichia coli BL21 DE3. Bacteria were grown in LB with Kanamycin (50 mg/ml) at 37 °C overnight. The culturing solution was inducted with 0.5 mM isopropyl 1-thio-β-D-galactopyranoside (IPTG) at optical density of 600nm (OD600) ∼0.6. Cells were centrifuged at RPM 8000 for 30 mins at 4 °C. The pellets were suspended in lysis buffer (20 mM MES. Na, 150 mM NaCl, 1 mM EDTA, 0.2 mM MgCl_2_, 5 mM DTT, protease inhibitor, pH 6.8) and sonicated on ice. The lysates were boiled in water for 20-30 min and centrifuged at RPM 55000, 4 °C for 50 min. The supernatant was dialyzed in dialysis buffer (20 mM MES. Na, 50 mM NaCl, 1 mM EDTA, 2 mM MgCl_2_, 2 mM DTT, pH 6.8) at 4 °C overnight. The protein was purified by cation-exchange chromatography (HiTrap™ SP Sepharose™ FF columns 5 mL) and size-exclusion chromatography (Superdex™ 75 Increase 10/300 GL). All proteins were buffer exchanged into 25LJmM Hepes, 150 mM NaCl, pH 7.4 by ultrafiltration (Amicon Ultra-0.5 mL 10K). Protein purity was verified by SDS-PAGE and concentrations were measured using NanoDrop. The protein was aliquoted and stored at −80LJ°C until use.

### Micropipette aspiration

In vitro MPA experiments were performed on a Nikon Ti2-A inverted fluorescence microscope (Nikon, Japan) equipped with a motorized stage, two motorized four-axis micromanipulators (PatchPro-5000, Scientifica), and a multi-trap optical tweezers system (Tweez305, Aresis, Slovenia). A long-pass emission filter (BLP02-561R-32, IDEX) was used to block wavelengths below 561 nm from the brightfield illumination. Micropipettes were fabricated from glass capillaries using a pipette puller (PUL-1000, World Precision Instruments). The tip was cut to achieve an opening diameter of 2–5 μm and then bent to an angle of ∼40° using a microforge (DMF1000, World Precision Instruments, as shown in Fig. S1A). The micropipette was backfilled with the synapsin buffer [25 mM Tris-HCl, 150 mM NaCl, and 0.5 mM TCEP (pH 7.4)] using a MICROFIL needle (World Precision Instruments). The pipette was mounted onto a micromanipulator and connected to an automated pressure controller (Flow-EZ, Fluigent; 1 Pa resolution). Experiments were conducted in glass-bottom dishes (D35-20-1.5-N, Cellvis). Prior to MPA, optical tweezers-assisted fusion was performed to generate large condensates (>5 μm) for improved measurement accuracy. Sample evaporation and air exchange were minimized by adding 1.5 mL of Milli-Q water to the edge of the dish and covering the dish and pipettes with a thin plastic wrap (Stretch-tite, Kirkland).

As the condensates in this study were observed to consistently wet the inner wall of uncoated micropipettes in vitro, the analysis followed the method described previously ^16^. Briefly, when the protein condensate wets the micropipette, purely viscous deformation follows 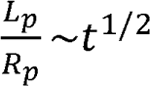 during MPA. In Figure 1c, the normalized deformation is 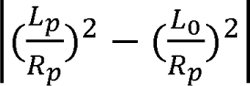 and the normalized time is 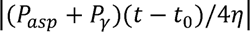. During brightfield imaging, the deformation is linear in time; with blue light excitation, however, the deformation of RGG-EGFP-RGG condensates gradually deviates from Newtonian behavior

### MAPAC experiments

Micropipettes for MAPAC were pulled into a cylindrical tip with an opening diameter of 0.5–1.0 μm (PUL-1000, World Precision Instruments). The tip was subsequently bent to an angle of ∼40° using a microforge (DMF1000, World Precision Instruments). Pipette resistance typically ranged from 25–65 MΩ. The micropipette was filled with an intracellular buffer (pH 7.2; osmolarity 270–290 mOsmol/kg). The intracellular buffer contained (in mM): 126 K-gluconate, 4 KCl, 10 Hepes, 2 Mg-ATP, 0.3 Na₂-GTP, and 10 phosphocreatine. All chemicals were sourced from Sigma-Aldrich. The filled micropipette was mounted to a headstage connected to an Axon 700B amplifier (Molecular Devices). A homemade pressure recording device utilizing an Arduino UNO R3 (ELEGOO) and a pressure sensor (FTVOGUE, Cat. No. B07N8SX347, 100 Pa resolution) was used to control aspiration pressure. Voltage recordings were processed through a 2-kHz filter, digitized at 10 kHz, and collected using Clampex 10.2 software (Molecular Devices). Liquid junction potentials were not compensated.

HEK 293T cells expressing fluorescently labeled condensates were targeted for whole-cell patch-clamp. The process began in voltage clamp mode. A small ejection pressure (∼−1 kPa) was applied until the pipette tip reached the targeted cell. Suction pressure was then applied and maintained to establish a giga seal near the targeted condensate. After giga seal formation, the recording was switched to current clamp mode (0 current). Sudden suction pressure pulses (>10 kPa) were applied to rupture the membrane and achieve the whole-cell configuration. The resting membrane voltage was recorded immediately. Once the whole-cell configuration was stable, suction pressures were applied to draw the nearby condensate into the pipette tip for subsequent measurement of its material properties.

### Cell culture

HEK 293T cells (ATCC) and HeLa cells (ATCC) were maintained in Dulbecco’s Modified Eagle’s Medium (DMEM) (11995065, Thermo Fisher Scientific) supplemented with 10% fetal bovine serum (FBS) (FB12999102, Fisherbrand) and 1% penicillin-streptomycin (15140122, Thermo Fisher Scientific). Cells were cultured at 37°C in a humidified incubator with 5–10% CO₂. For routine passaging, approximately 1.0 x 10 cells were seeded into 100-mm plastic dishes.

### Transfection and labeling

3.0 x 10^5^ cells were seeded onto 35-mm glass bottom dishes (Cat. No. D35-20-1.5-N, Cellvis) that had been pre-coated with Matrigel Matrix (Cat. No. 47743-715-EA, Corning Life Sciences). Twenty-four hours after seeding, cells were transiently transfected using Lipofectamine P3000 reagent (Thermo Fisher Scientific). The transfection mixture contained 250 μL Opti-MEM, 2 μL P3000 reagent, 7.5 μL Lipofectamine reagent, 1 μg plasmid. The plasmid pEGFP-C1-G3BP1-WT (Addgene, 135997) and pRK5_mEGFP-NPM1(Addgene, 237667) are ordered from Addgene, and the plasmid mCherry-G3BP1 is a gift from Prof. Clifford P Brangwynne lab. Cellular ROS levels were measured using a ROS Detection Assay Kit (Abcam, ab186029) according to the manufacturer’s protocol.

### FRAP

FRAP experiments were performed on a total internal reflection fluorescence microscope (DMi8 TIRF, Leica, Germany) equipped with an Infinity Scanner system (Leica, Germany). All images were acquired using a 100X oil objective at 1 Hz. The FRAP experiments were carried out using the protocol as described in reference^42^.

### Light power density measurement

The light power was measured using an optical power meter (843-R, Newport), and the reported values are the maximum of at least three independent measurements. For fluorescence illumination, area of the light was quantified by acquiring an image of the illuminated region when it fully fits within the field of view; when the exposure exceeded the imaging area, the photobleached region after 5 minutes of illumination was used to estimate the light area. For brightfield illumination, the illuminated area was assumed to be equivalent to the detector size (1 cm^2^). The light power density was then calculated by dividing the measured light power by the corresponding illuminated area.

### UV–vis spectroscopy

UV–vis absorption spectra were recorded using a NanoDrop spectrophotometer (Thermo Fisher Scientific). After exposure to blue light for different durations, an equal volume of 8 M urea was added to the sample to dissolve the phase-separated droplets and generate a homogeneous solution. For each measurement, 1-2 μl of sample was loaded directly onto the pedestal, and spectra were collected over the wavelength range of 220–750 nm. After blank subtraction, the absorbance spectrum of each sample was analyzed using the instrument software.

### SDS-PAGE

SDS-PAGE gels in Figure 1h and 3i were run using NuPAGE 4–12% Bis-Tris gels (Invitrogen). The gel in Figure 2c was run using Mini-PROTEAN TGA gels. The gels are visualized with GelCode Blue Safe Protein Coomassie stain (Thermo Scientific).

## References

1. Shin, Y. & Brangwynne, C. P. Liquid phase condensation in cell physiology and disease. Science 357, eaaf4382 (2017).

2. Alberti, S., Gladfelter, A. & Mittag, T. Considerations and challenges in studying liquid-liquid phase separation and biomolecular condensates. Cell 176, 419–434 (2019).

3. Banani, S. F., Lee, H. O., Hyman, A. A. & Rosen, M. K. Biomolecular condensates: organizers of cellular biochemistry. Nature reviews Molecular cell biology 18, 285–298 (2017).

4. Stroberg, W. & Schnell, S. Do cellular condensates accelerate biochemical reactions? Lessons from microdroplet chemistry. Biophys. J. 115, 3–8 (2018).

5. Gil-Garcia, M., et al. Local environment in biomolecular condensates modulates enzymatic activity across length scales. Nature communications 15, 3322 (2024).

6. Nott, T. J., Craggs, T. D. & Baldwin, A. J. Membraneless organelles can melt nucleic acid duplexes and act as biomolecular filters. Nature chemistry 8, 569–575 (2016).

7. Dai, Y., et al. Interface of biomolecular condensates modulates redox reactions. Chem 9, 1594–1609 (2023).

8. Dumelie, J. G., et al. Biomolecular condensates create phospholipid-enriched microenvironments. Nature Chemical Biology 20, 302–313 (2024).

9. Ambadi Thody, S., et al. Small-molecule properties define partitioning into biomolecular condensates. Nature Chemistry 16, 1794–1802 (2024).

10. Holehouse, A. S. & Alberti, S. Molecular determinants of condensate composition. Mol. Cell 85, 290–308 (2025).

11. Kilgore, H. R., et al. Distinct chemical environments in biomolecular condensates. Nature Chemical Biology 20, 291–301 (2024).

12. Rekhi, S. & Mittal, J. Biomolecular Condensates Act as Distinct Solvation Environments that Reshape Amino Acid pKLJ Values. bioRxiv, 2026.01. 11.698869 (2026).

13. Lyon, A. S., Peeples, W. B. & Rosen, M. K. A framework for understanding the functions of biomolecular condensates across scales. Nature reviews Molecular cell biology 22, 215–235 (2021).

14. Kroschwald, S., et al. Different material states of Pub1 condensates define distinct modes of stress adaptation and recovery. Cell reports 23, 3327–3339 (2018).

15. Galvanetto, N., et al. Material properties of biomolecular condensates emerge from nanoscale dynamics. Proceedings of the National Academy of Sciences 122, e2424135122 (2025).

16. Wang, H., et al. Live-cell quantification reveals viscoelastic regulation of synapsin condensates by α-synuclein. Science Advances 11, eads7627 (2025).

17. Tejedor, A. R., Collepardo-Guevara, R., Ramírez, J. & Espinosa, J. R. Time-dependent material properties of aging biomolecular condensates from different viscoelasticity measurements in molecular dynamics simulations. The Journal of Physical Chemistry B 127, 4441–4459 (2023).

18. Jawerth, L., et al. Protein condensates as aging Maxwell fluids. Science 370, 1317 (2020).

19. Emmanouilidis, L., et al. A solid beta-sheet structure is formed at the surface of FUS droplets during aging. Nature Chemical Biology 20, 1044–1052 (2024).

20. Yu, W., et al. Aging-dependent evolving electrochemical potentials of biomolecular condensates regulate their physicochemical activities. Nature Chemistry, 1–11 (2025).

21. Wang, J., et al. Phase separation of the nuclear pore complex facilitates selective nuclear transport to regulate plant defense against pathogen and pest invasion. Molecular Plant 16, 1016–1030 (2023).

22. Li, Z., et al. Phase-separated NDF− FACT condensates facilitate transcription elongation on chromatin. Nat. Cell Biol., 1–14 (2025).

23. Guo, P., et al. PI4P-mediated solid-like Merlin condensates orchestrate Hippo pathway regulation. Science 385, eadf4478 (2024).

24. Nott, T. J., et al. Phase transition of a disordered nuage protein generates environmentally responsive membraneless organelles. Mol. Cell 57, 936–947 (2015).

25. Linsenmeier, M., et al. Dynamic arrest and aging of biomolecular condensates are modulated by low-complexity domains, RNA and biochemical activity. Nature Communications 13, 3030 (2022).

26. Aguzzi, A. & Altmeyer, M. Phase separation: linking cellular compartmentalization to disease. Trends Cell Biol. 26, 547–558 (2016).

27. Alberti, S. & Dormann, D. Liquid–liquid phase separation in disease. Annu. Rev. Genet. 53, 171–194 (2019).

28. Wang, H. & Shi, Z. The rheology and interfacial properties of biomolecular condensates. Biophysical Reviews, 1–25 (2025).

29. Shen, Y., et al. The liquid-to-solid transition of FUS is promoted by the condensate surface. Proceedings of the National Academy of Sciences 120, e2301366120 (2023).

30. Murakami, T., et al. ALS/FTD mutation-induced phase transition of FUS liquid droplets and reversible hydrogels into irreversible hydrogels impairs RNP granule function. Neuron 88, 678–690 (2015).

31. Taniue, K. & Akimitsu, N. Aberrant phase separation and cancer. The FEBS Journal 289, 17 (2021).

32. Patel, A., et al. A liquid-to-solid phase transition of the ALS protein FUS accelerated by disease mutation. Cell 162, 1066–1077 (2015).

33. Tinevez, J. et al. in Methods in enzymology 291–309 (Elsevier, 2012).

34. Icha, J., Weber, M., Waters, J. C. & Norden, C. Phototoxicity in live fluorescence microscopy, and how to avoid it. Bioessays 39, 1700003 (2017).

35. Swoboda, M., et al. Enzymatic oxygen scavenging for photostability without pH drop in single-molecule experiments. ACS nano 6, 6364–6369 (2012).

36. Demchenko, A. P. Photobleaching of organic fluorophores: quantitative characterization, mechanisms, protection. Methods and applications in fluorescence 8, 022001 (2020).

37. Schieber, M. & Chandel, N. S. ROS function in redox signaling and oxidative stress. Current biology 24, R453–R462 (2014).

38. Lennicke, C. & Cochemé, H. M. Redox metabolism: ROS as specific molecular regulators of cell signaling and function. Mol. Cell 81, 3691–3707 (2021).

39. Murphy, M. P., et al. Guidelines for measuring reactive oxygen species and oxidative damage in cells and in vivo. Nature metabolism 4, 651–662 (2022).

40. Liguori, I., et al. Oxidative stress, aging, and diseases. Clinical interventions in aging, 757–772 (2018).

41. Schuster, B. S., et al. Controllable protein phase separation and modular recruitment to form responsive membraneless organelles. Nature communications 9, 2985 (2018).

42. Wang, H., Kelley, F. M., Milovanovic, D., Schuster, B. S. & Shi, Z. Surface tension and viscosity of protein condensates quantified by micropipette aspiration. Biophysical Reports 1 (2021).

43. Roggeveen, J. V., Wang, H., Shi, Z. & Stone, H. A. A calibration-free model of micropipette aspiration for measuring properties of protein condensates. Biophys. J. 123, 1393–1403 (2024).

44. Guevorkian, K., Colbert, M., Durth, M., Dufour, S. & Brochard-Wyart, F. Aspiration of biological viscoelastic drops. Phys. Rev. Lett. 104, 218101 (2010).

45. Shi, Z., Graber, Z. T., Baumgart, T., Stone, H. A. & Cohen, A. E. Cell membranes resist flow. Cell 175, 1769–1779. e13 (2018).

46. Hirano, M., et al. A highly photostable and bright green fluorescent protein. Nat. Biotechnol. 40, 1132–1142 (2022).

47. Flory, P. J. Viscosities of linear polyesters. An exact relationship between viscosity and chain length. J. Am. Chem. Soc. 62, 1057–1070 (1940).

48. Park, A. S., Ding, E. A. & Schuster, B. S. POMPOMS: Crosslinked biomolecular condensates as a versatile platform for multifunctional protein microparticles. Biomacromolecules 26, 7605–7620 (2025).

49. Bulina, M. E., et al. A genetically encoded photosensitizer. Nat. Biotechnol. 24, 95–99 (2006).

50. Shu, X., et al. A genetically encoded tag for correlated light and electron microscopy of intact cells, tissues, and organisms. PLoS biology 9, e1001041 (2011).

51. Milovanovic, D., Wu, Y., Bian, X. & De Camilli, P. A liquid phase of synapsin and lipid vesicles. Science 361, 604–607 (2018).

52. Hoffmann, C., et al. Synapsin condensation controls synaptic vesicle sequestering and dynamics. Nature Communications 14, 6730 (2023).

53. Zbinden, A., Pérez-Berlanga, M., De Rossi, P. & Polymenidou, M. Phase separation and neurodegenerative diseases: a disturbance in the force. Developmental cell 55, 45–68 (2020).

54. Mathieu, C., Pappu, R. V. & Taylor, J. P. Beyond aggregation: Pathological phase transitions in neurodegenerative disease. Science 370, 56–60 (2020).

55. Ray, S., et al. α-Synuclein aggregation nucleates through liquid–liquid phase separation. Nature chemistry 12, 705–716 (2020).

56. Kanaan, N. M., Hamel, C., Grabinski, T. & Combs, B. Liquid-liquid phase separation induces pathogenic tau conformations in vitro. Nature communications 11, 2809 (2020).

57. Ettinger, A. & Wittmann, T. Fluorescence live cell imaging. Methods Cell Biol. 123, 77–94 (2014).

58. Sharma, B., Baum, J. & Shi, Z. The material properties and crosstalk between tau and alpha-synuclein condensates. Biophys. J. 123, 71a (2024).

59. Zheng, Q., Jockusch, S., Zhou, Z. & Blanchard, S. C. The contribution of reactive oxygen species to the photobleaching of organic fluorophores. Photochem. Photobiol. 90, 448–454 (2014).

60. Kelley, F. M., et al. Controlled and orthogonal partitioning of large particles into biomolecular condensates. Nature communications 16, 3521 (2025).

61. Taylor, N. O., Wei, M., Stone, H. A. & Brangwynne, C. P. Quantifying dynamics in phase-separated condensates using fluorescence recovery after photobleaching. Biophys. J. 117, 1285–1300 (2019).

62. Spicer, C. D. & Davis, B. G. Selective chemical protein modification. Nature communications 5, 4740 (2014).

63. Konigsberg, W. in Methods in enzymology 185–188 (Elsevier, 1972).

64. Milligan, J. R., et al. Repair of oxidative DNA damage by amino acids. Nucleic Acids Res. 31, 6258–6263 (2003).

65. Heinecke, J. W., Li, W., Daehnke 3rd, H. L. & Goldstein, J. A. Dityrosine, a specific marker of oxidation, is synthesized by the myeloperoxidase-hydrogen peroxide system of human neutrophils and macrophages. J. Biol. Chem. 268, 4069–4077 (1993).

66. Yu, W., et al. Origins of the Intrinsic Redox Activity of Biomolecular Condensates. J. Am. Chem. Soc. (2026).

67. Imlay, J. A., Chin, S. M. & Linn, S. Toxic DNA damage by hydrogen peroxide through the Fenton reaction in vivo and in vitro. Science 240, 640–642 (1988).

68. Pawlik, K., Ostrowska, M. & Gumienna-Kontecka, E. Systematic Model Peptide Studies: A Crucial Step To Understand the Coordination Chemistry of Mn (II) and Fe (II) in Proteins. Inorg. Chem. 64, 5472–5486 (2025).

69. Miao, R., et al. Enhanced volatile fatty acids production from anaerobic fermentation of grease trap waste by Fenton pretreatment. Water Cycle 6, 278–284 (2025).

70. Qu, X., Song, W. & Shen, Z. A highly selective NIR fluorescent turn-on probe for hydroxyl radical and its application in living cell images. Frontiers in Chemistry 7, 598 (2019).

71. Protter, D. S. & Parker, R. Principles and properties of stress granules. Trends Cell Biol. 26, 668–679 (2016).

72. Jomova, K., et al. Several lines of antioxidant defense against oxidative stress: antioxidant enzymes, nanomaterials with multiple enzyme-mimicking activities, and low-molecular-weight antioxidants. Arch. Toxicol. 98, 1323–1367 (2024).

73. Glauninger, H., et al. Transcriptome-wide mRNP condensation precedes stress granule formation and excludes new mRNAs. Mol. Cell 85, 4393–4409. e11 (2025).

74. Mohammadiaria, M. ROS-induced voltage-gated ion channel expression and electrophysiological remodeling in malignant human cells. npj Systems Biology and Applications 11, 119 (2025).

75. Ighodaro, O. M. & Akinloye, O. A. First line defence antioxidants-superoxide dismutase (SOD), catalase (CAT) and glutathione peroxidase (GPX): Their fundamental role in the entire antioxidant defence grid. Alexandria journal of medicine 54, 287–293 (2018).

76. Feric, M., et al. Coexisting liquid phases underlie nucleolar subcompartments. Cell 165, 1686–1697 (2016).

77. Garg, A., Brasnett, C., Marrink, S. J., Koren, K. & Kjaergaard, M. Oxygen depletion in biomolecular condensates is dominated by macromolecular density. Nature Communications (2026).

78. Ibrahim, K. A., Naidu, A. S., Miljkovic, H., Radenovic, A. & Yang, W. Label-free techniques for probing biomolecular condensates. ACS nano 18, 10738–10757 (2024).

79. Dörner, K., et al. Fluorescent protein and peptide tags alter condensate formation and dynamics in vivo and in vitro. EMBO Rep. 27, 89 (2025).

80. Branon, T. C., et al. Efficient proximity labeling in living cells and organisms with TurboID. Nat. Biotechnol. 36, 880–887 (2018).

81. Qin, W., Cho, K. F., Cavanagh, P. E. & Ting, A. Y. Deciphering molecular interactions by proximity labeling. Nature methods 18, 133–143 (2021).

82. Kim, S., et al. Hidden route of protein damage through oxygen-confined photooxidation. Nature Communications 15, 10873 (2024).

83. Sheikhhassani, V., Wong, F. H., Bonn, D., Schmit, J. D. & Mashaghi, A. Optically driven control of mechanochemistry and fusion dynamics of biomolecular condensates via thymine dimerization. Nature Communications (2026).

