## Supplementary Information for "Biomolecular condensates provide a unique environment for redox-mediated protein crosslinking"

**This supplement contains:**

Table S1

Figure S1 – S8

Supplementary Discussion

**Table S1:** Sequence information of purified proteins used in this study, with the fluorescent protein highlighted in green (for EGFP, miniSOG, cfGFP) or red (for KillerRed). For Tau, binding sites for AF488 maleimide are highlighted in yellow.

| **Protein** | **Sequence** |
| --- | --- |
| RGG-RGG | MESNQSNNGGSGNAALNRGGRYVPPHLRGGDGGAAAAASAGGDDRRGGAGGGGYRRGGGNSGGGGGGGYDRGYNDNRDDRDNRGGSGGYGRDRNYEDRGYNGGGGGGGNRGYNNNRGGGGGGYNRQDRGDGGSSNFSRGGYNNRDEGSDNRGSGRSYNNDRRDNGGDGEFGKLMESNQSNNGGSGNAALNRGGRYVPPHLRGGDGGAAAAASAGGDDRRGGAGGGGYRRGGGNSGGGGGGGYDRGYNDNRDDRDNRGGSGGYGRDRNYEDRGYNGGGGGGGNRGYNNNRGGGGGGYNRQDRGDGGSSNFSRGGYNNRDEGSDNRGSGRSYNNDRRDNGGDGLEHHHHHH |
| RGG-EGFP-RGG | MESNQSNNGGSGNAALNRGGRYVPPHLRGGDGGAAAAASAGGDDRRGGAGGGGYRRGGGNSGGGGGGGYDRGYNDNRDDRDNRGGSGGYGRDRNYEDRGYNGGGGGGGNRGYNNNRGGGGGGYNRQDRGDGGSSNFSRGGYNNRDEGSDNRGSGRSYNNDRRDNGGDGGETVRFQSMVSKGEELFTGVVPILVELDGDVNGHKFSVSGEGEGDATYGKLTLKFICTTGKLPVPWPTLVTTLTYGVQCFSRYPDHMKQHDFFKSAMPEGYVQERTIFFKDDGNYKTRAEVKFEGDTLVNRIELKGIDFKEDGNILGHKLEYNYNSHNVYIMADKQKNGIKVNFKIRHNIEDGSVQLADHYQQNTPIGDGPVLLPDNHYLSTQSKLSKDPNEKRDHMVLLEFVTAAGITLGMDELYKGGGSENLYFQGEFGKLMESNQSNNGGSGNAALNRGGRYVPPHLRGGDGGAAAAASAGGDDRRGGAGGGGYRRGGGNSGGGGGGGYDRGYNDNRDDRDNRGGSGGYGRDRNYEDRGYNGGGGGGGNRGYNNNRGGGGGGYNRQDRGDGGSSNFSRGGYNNRDEGSDNRGSGRSYNNDRRDNGGDGLEHHHHHH |
| RGG-cfGFP-RGG | MESNQSNNGGSGNAALNRGGRYVPPHLRGGDGGAAAAASAGGDDRRGGAGGGGYRRGGGNSGGGGGGGYDRGYNDNRDDRDNRGGSGGYGRDRNYEDRGYNGGGGGGGNRGYNNNRGGGGGGYNRQDRGDGGSSNFSRGGYNNRDEGSDNRGSGRSYNNDRRDNGGDGGETVRFQSMVSKGEELFTGVVPILVELDGDVNGHKFSVSGEGEGDATYGKLTLKFISTTGKLPVPWPTLVTTLTYGVQMFARYPDHMKQHDFFKSAMPEGYVQERTIFFKDDGNYKTRAEVKFEGDTLVNRIELKGIDFKEDGNILGHKLEYNYNSHNVYITADKQKNGIKANFKIRHNIEDGGVQLADHYQQNTPIGDGPVLLPDNHYLSTQSKLSKDPNEKRDHMVLLEFVTAAGITLGMDELYKGGGSENLYFQGEFGKLMESNQSNNGGSGNAALNRGGRYVPPHLRGGDGGAAAAASAGGDDRRGGAGGGGYRRGGGNSGGGGGGGYDRGYNDNRDDRDNRGGSGGYGRDRNYEDRGYNGGGGGGGNRGYNNNRGGGGGGYNRQDRGDGGSSNFSRGGYNNRDEGSDNRGSGRSYNNDRRDNGGDGLEHHHHHH |
| RGG-KillerRed-RGG | MESNQSNNGGSGNAALNRGGRYVPPHLRGGDGGAAAAASAGGDDRRGGAGGGGYRRGGGNSGGGGGGGYDRGYNDNRDDRDNRGGSGGYGRDRNYEDRGYNGGGGGGGNRGYNNNRGGGGGGYNRQDRGDGGSSNFSRGGYNNRDEGSDNRGSGRSYNNDRRDNGGDGGETVRFQSMGSEGGPALFQSDMTFKIFIDGEVNGQKFTIVADGSSKFPHGDFNVHAVCETGKLPMSWKPICHLIQYGEPFFARYPDGISHFAQECFPEGLSIDRTVRFENDGTMTSHHTYELDDTCVVSRITVNCDGFQPDGPIMRDQLVDILPNETHMFPHGPNAVRQLAFIGFTTADGGLMMGHFDSKMTFNGSRAIEIPGPHFVTIITKQMRDTSDKRDHVCQREVAYAHSVPRITSAIGSDEDGGGSENLYFQGEFGKLMESNQSNNGGSGNAALNRGGRYVPPHLRGGDGGAAAAASAGGDDRRGGAGGGGYRRGGGNSGGGGGGGYDRGYNDNRDDRDNRGGSGGYGRDRNYEDRGYNGGGGGGGNRGYNNNRGGGGGGYNRQDRGDGGSSNFSRGGYNNRDEGSDNRGSGRSYNNDRRDNGGDGLEHHHHHH |
| RGG-MiniSOG-RGG | MESNQSNNGGSGNAALNRGGRYVPPHLRGGDGGAAAAASAGGDDRRGGAGGGGYRRGGGNSGGGGGGGYDRGYNDNRDDRDNRGGSGGYGRDRNYEDRGYNGGGGGGGNRGYNNNRGGGGGGYNRQDRGDGGSSNFSRGGYNNRDEGSDNRGSGRSYNNDRRDNGGDGGETVRFQSMEKSFVITDPRLPDNPIIFASDGFLELTEYSREEILGRNGRFLQGPETDQATVQKIRDAIRDQREITVQLINYTKSGKKFWNLLHLQPMRDQKGELQYFIGVQLDGGGGSENLYFQGEFGKLMESNQSNNGGSGNAALNRGGRYVPPHLRGGDGGAAAAASAGGDDRRGGAGGGGYRRGGGNSGGGGGGGYDRGYNDNRDDRDNRGGSGGYGRDRNYEDRGYNGGGGGGGNRGYNNNRGGGGGGYNRQDRGDGGSSNFSRGGYNNRDEGSDNRGSGRSYNNDRRDNGGDGLEHHHHHH |
| Tau | MAEPRQEFEVMEDHAGTYGLGDRKDQGGYTMHQDQEGDTDAGLKESPLQTPTEDGSEEPGSETSDAKSTPTAEDVTAPLVDEGAPGKQAAAQPHTEIPEGTTAEEAGIGDTPSLEDEAAGHVTQARMVSKSKDGTGSDDKKAKGADGKTKIATPRGAAPPGQKGQANATRIPAKTPPAPKTPPSSGEPPKSGDRSGYSSPGSPGTPGSRSRTPSLPTPPTREPKKVAVVRTPPKSPSSAKSRLQTAPVPMPDLKNVKSKIGSTENLKHQPGGGKVQIINKKLDLSNVQSKCGSKDNIKHVPGGGSVQIVYKPVDLSKVTSKCGSLGNIHHKPGGGQVEVKSEKLDFKDRVQSKIGSLDNITHVPGGGNKKIETHKLTFRENAKAKTDHGAEIVYKSPVVSGDTSPRHLSNVSSTGSIDMVDSPQLATLADEVSASLAKQGL |
| EGFP-Synapsin | MVSKGEELFTGVVPILVELDGDVNGHKFSVSGEGEGDATYGKLTLKFICTTGKLPVPWPTLVTTLTYGVQCFSRYPDHMKQHDFFKSAMPEGYVQERTIFFKDDGNYKTRAEVKFEGDTLVNRIELKGIDFKEDGNILGHKLEYNYNSHNVYIMADKQKNGIKVNFKIRHNIEDGSVQLADHYQQNTPIGDGPVLLPDNHYLSTQSALSKDPNEKRDHMVLLEFVTAAGITLGMDELYKSGLRSMNYLRRRLSDSNFMANLPNGYMTDLQRPQPPPPPPGAHSPGATPGPGTATAERSSGVAPAASPAAPSPGSSGGGGFFSSLSNAVKQTTAAAAATFSEQVGGGSGGAGRGGAASRVLLVIDEPHTDWAKYFKGKKIHGEIDIKVEQAEFSDLNLVAHANGGFSVDMEVLRNGVKVVRSLKPDFVLIRQHAFSMARNGDYRSLVIGLQYAGIPSVNSLHSVYNFCDKPWVFAQMVRLHKKLGTEEFPLIDQTFYPNHKEMLSSTTYPVVVKMGHAHSGMGKVKVDNQHDFQDIASVVALTKTYATAEPFIDAKYDVRVQKIGQNYKAYMRTSVSGNWKTNTGSAMLEQIAMSDRYKLWVDTCSEIFGGLDICAVEALHGKDGRDHIIEVVGSSMPLIGDHQDEDKQLIVELVVNKMAQALPRQRQRDASPGRGSHGQTPSPGALPLGRQTSQQPAGPPAQQRPPPQGGPPQPGPGPQRQGPPLQQRPPPQGQQHLSGLGPPAGSPLPQRLPSPTSAPQQPASQAAPPTQGQGRQSRPVAGGPGAPPAARPPASPSPQRQAGPPQATRQTSVSGPAPPKASGAPPGGQQRQGPPQKPPGPAGPTRQASQAGPVPRTGPPTTQQPRPSGPGPAGRPKPQLAQKPSQDVPPPATAAAGGPPHPQLNKSQSLTNAFNLPEPAPPRPSLSQDEVKAETIRSLRKSFASLFSD |
| EGFP-Synapsin IDR | MVSKGEELFTGVVPILVELDGDVNGHKFSVSGEGEGDATYGKLTLKFICTTGKLPVPWPTLVTTLTYGVQCFSRYPDHMKQHDFFKSAMPEGYVQERTIFFKDDGNYKTRAEVKFEGDTLVNRIELKGIDFKEDGNILGHKLEYNYNSHNVYIMADKQKNGIKVNFKIRHNIEDGSVQLADHYQQNTPIGDGPVLLPDNHYLSTQSALSKDPNEKRDHMVLLEFVTAAGITLGMDELYKSGLRSQALPRQRQRDASPGRGSHGQTPSPGALPLGRQTSQQPAGPPAQQRPPPQGGPPQPGPGPQRQGPPLQQRPPPQGQQHLSGLGPPAGSPLPQRLPSPTSAPQQPASQAAPPTQGQGRQSRPVAGGPGAPPAARPPASPSPQRQAGPPQATRQTSVSGPAPPKASGAPPGGQQRQGPPQKPPGPAGPTRQASQAGPVPRTGPPTTQQPRPSGPGPAGRPKPQLAQKPSQDVPPPATAAAGGPPHPQLNKSQSLTNAFNLPEPAPPRPSLSQDEVKAETIRSLRKSFASLFSD |
| Alpha-Synuclein | MDVFMKGLSKAKEGVVAAAEKTKQGVAEAAGKTKEGVLYVGSKTKEGVVHGVATVAEKTKEQVTNVGGAVVTGVTAVAQKTVEGAGSIAAATGFVKKDQLGKNEEGAPQEGILEDMPVDPDNEAYEMPSEEGYQDYEPEA |

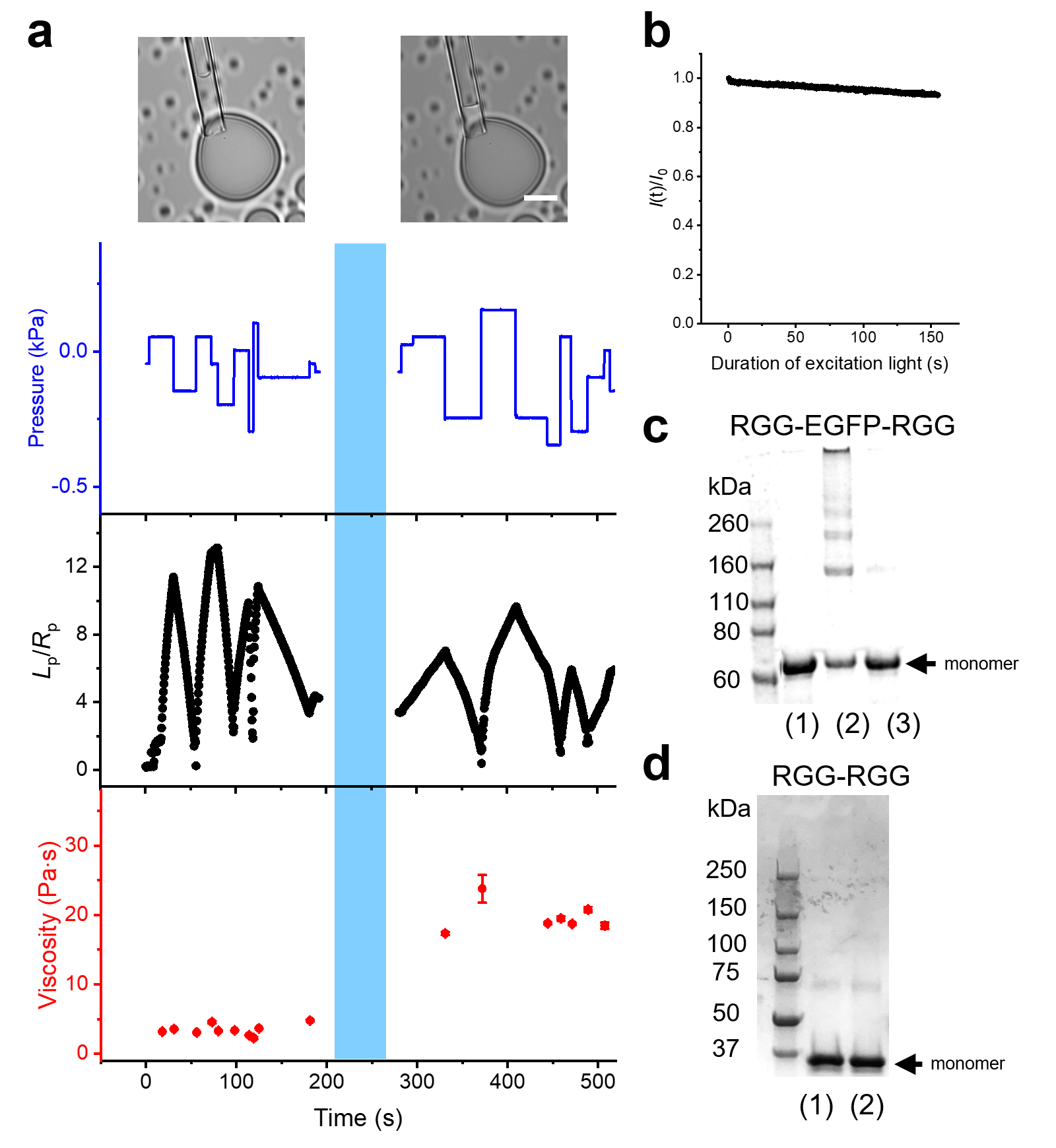

**Figure S1**: Blue light and the EGFP tag are essential for the solidification of RGG condensates. (**a**) Images (top) and MPA measurements (bottom) on a RGG-EGFP-RGG condensate under brightfield before and after 60 s blue light. Blue: aspiration pressure. Black: aspiration length (*L*_p_) normalized to the micropipette radius (*R*_p_). Red: viscosity calculated from every pressure step. Blue shade represents 60s 0.122 W/cm^2^ blue light. Scale bar: 10 μm. (**b**) Photobleaching curve of RGG-EGFP-RGG condensates under 0.122 W/cm^2^ blue light. (**c**) SDS–PAGE gel of RGG-EGFP-RGG aliquots under 3 different conditions: (1) no blue light + 23 °C, (2) ‘blue light’ + 23 °C, (3) ‘blue light’ + 42 °C. (**d**) SDS–PAGE gel RGG-RGG aliquots under 2 different conditions: (1) no blue light + 23 °C, (2) ‘blue light’ + 23 °C. All samples are 10 μM of protein in 150 mM NaCl, 20 mM Tris, pH 7.5. The samples are phase-separated under 23 °C and homogeneous under 42 °C. ‘Blue light’ refers to 15 min of 0.133 W/cm^2^ 480 nm LED exposure.

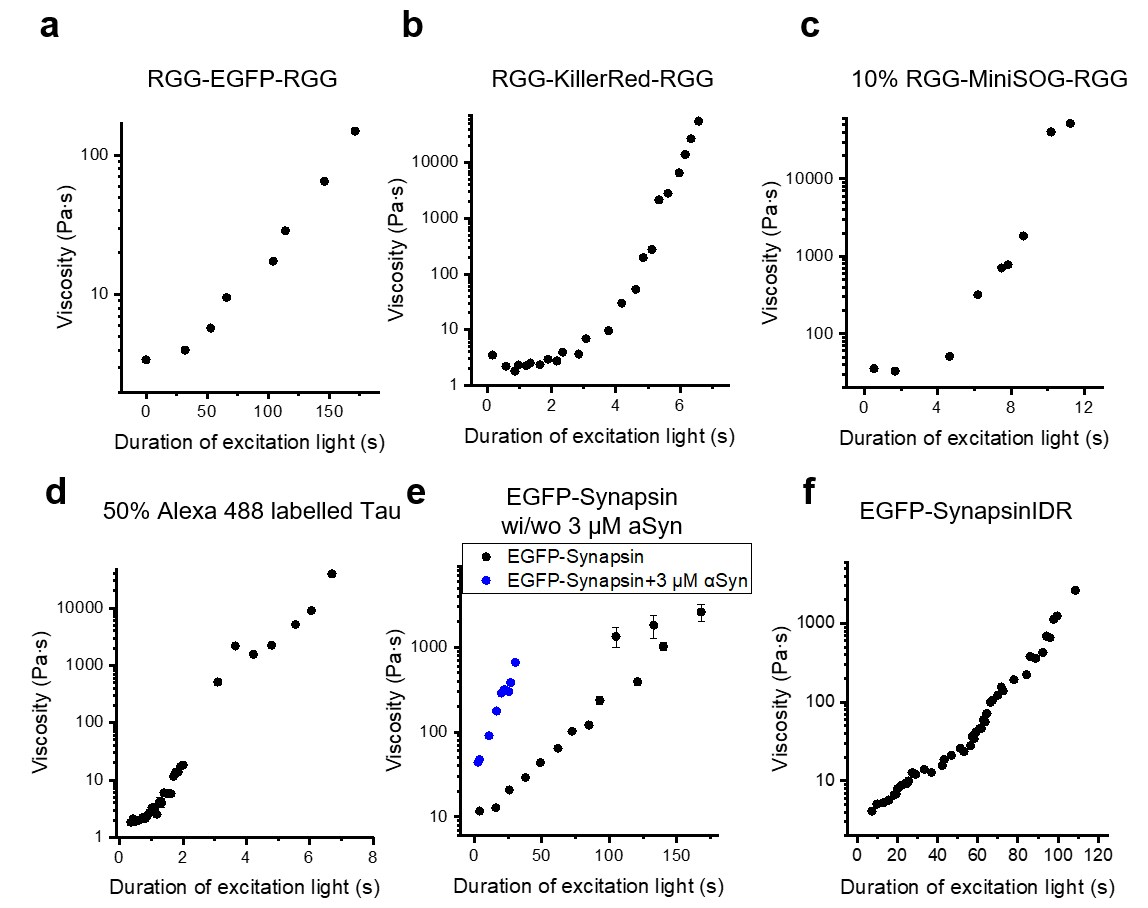

**Figure S2**: Representative MPA measurements of excitation-induced condensate viscosity increase for proteins listed in **Figure 2a**.

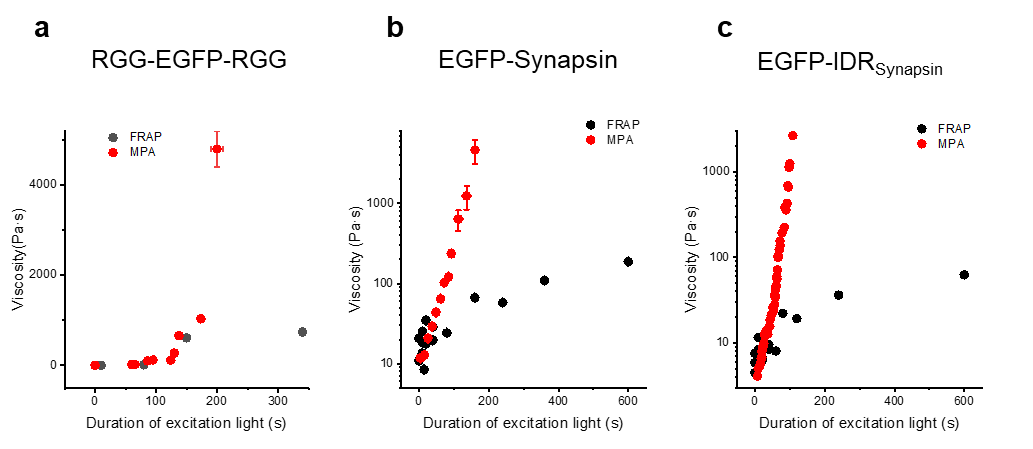

**Figure S3**: Comparison of condensate viscosities derived from MPA and FRAP.

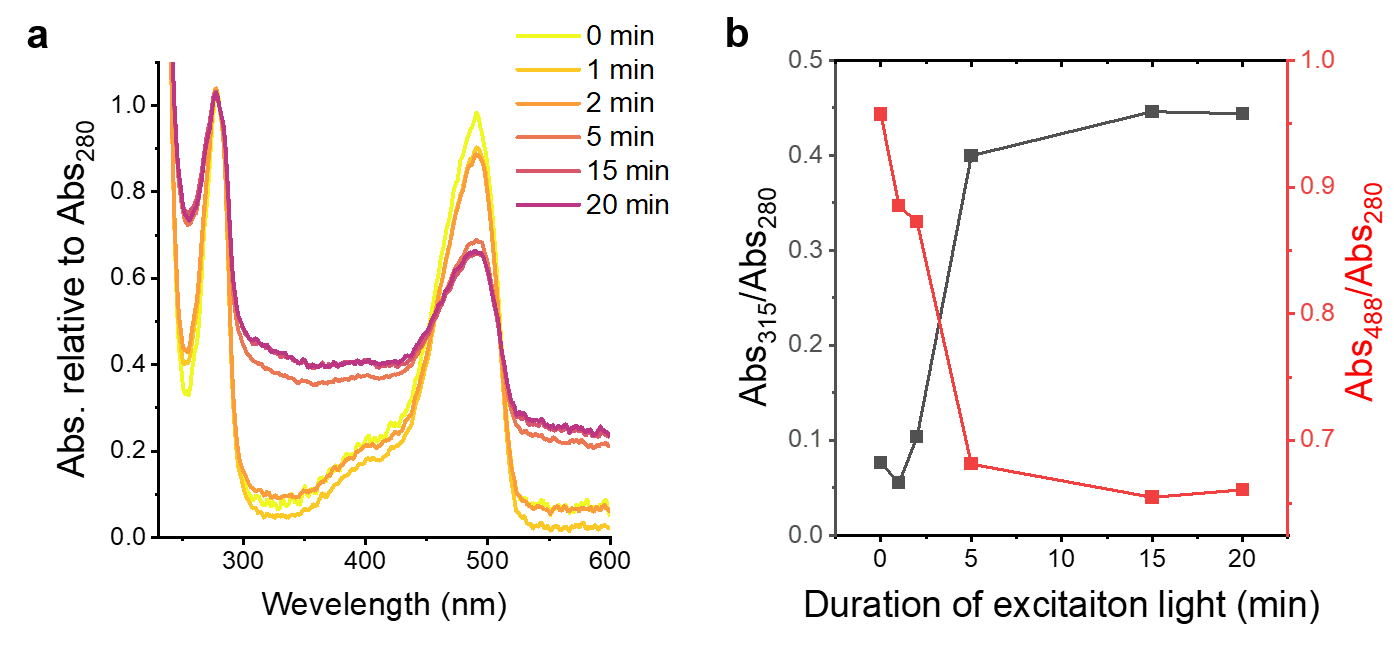

**Figure S4**: Compositional change during the excitation of RGG-EGFP-RGG condensates.

(**a**) Normalized UV-Vis absorbance spectra of RGG-EGFP-RGG with increasing durations of blue light exposure (0-20 min). The spectra were normalized to the protein’s absorbance at 280 nm. (**b**) Normalized absorbance at 315 nm (black) showing the formation of dityrosine. Normalized absorbance at 488 nm (red) mainly associated with the photobleaching of EGFP.

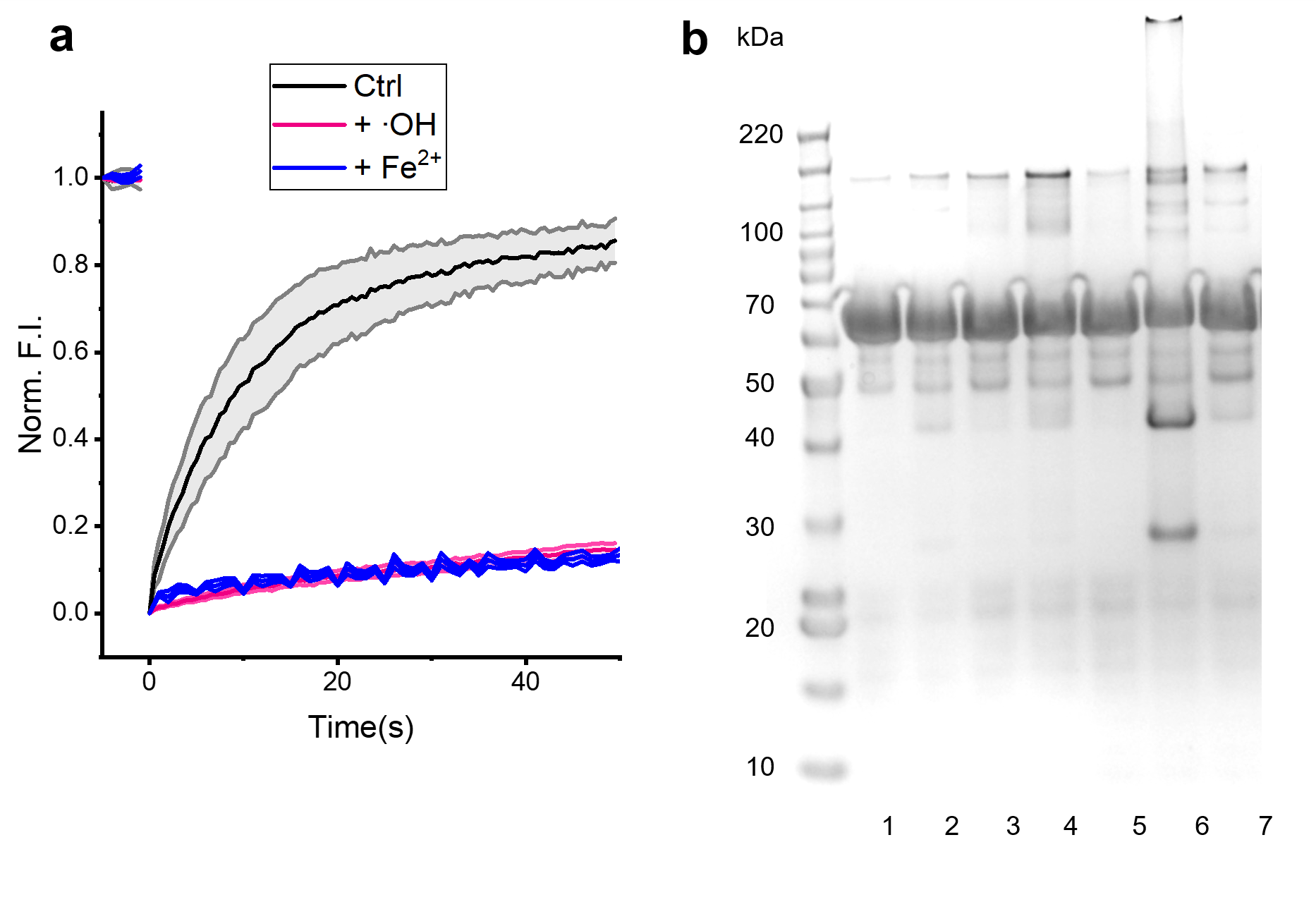

**Figure S5**: Full image of the gel in **Figure 4c**. Numbers on the left indicate the marker sizes(kDa). Numbers on the bottom label the number of samples; 1:no treatment; 2: +10 mM H_2_O_2_ @ 42 ℃; 3: +10 mM H_2_O_2_ @ 23 ℃; 4: +1 mM FeCl_2_ @ 42 ℃; 5: +1 mM FeCl_2_ @ 23 ℃; 6: +10 mM H_2_O_2_ and 1 mM FeCl_2_ @ 42 ℃; 7: +10 mM H_2_O_2_ and 1 mM FeCl_2_ @ 23 ℃.

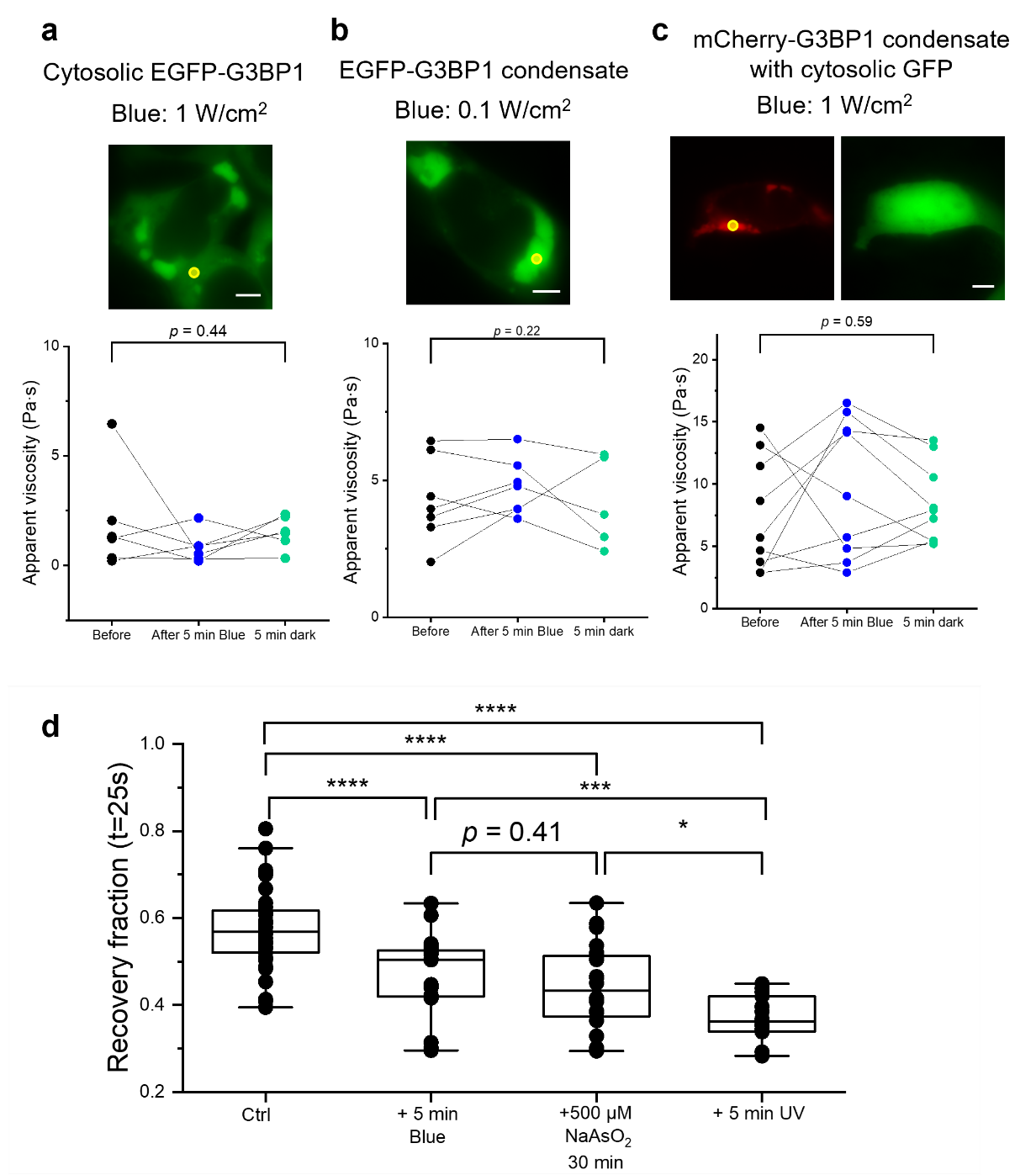

**Figure S6:** Comparison of different stress levels on the material properties of SGs

(**a**-**c**) Change of the apparent viscosity quantified by FRAP of before and after 5 min blue light and following 5 min recovery in the dark. a: 0.133 W/cm^2^ blue light for 5-min blue light excitation, bleaching spots are in condensate. (b): Bleaching spots are in cytosol, 0.974 W/cm^2^ blue light for 5-min illumination. (c): mCherry–G3BP1 condensates with cytosolic GFP, bleaching spots are in condensate, 0.974 W/cm^2^ blue light for 5-min illumination. (**d**-**e**) The recovery time and fraction quantified from FRAP on SGs in different conditions. Scale bar: 5 μm

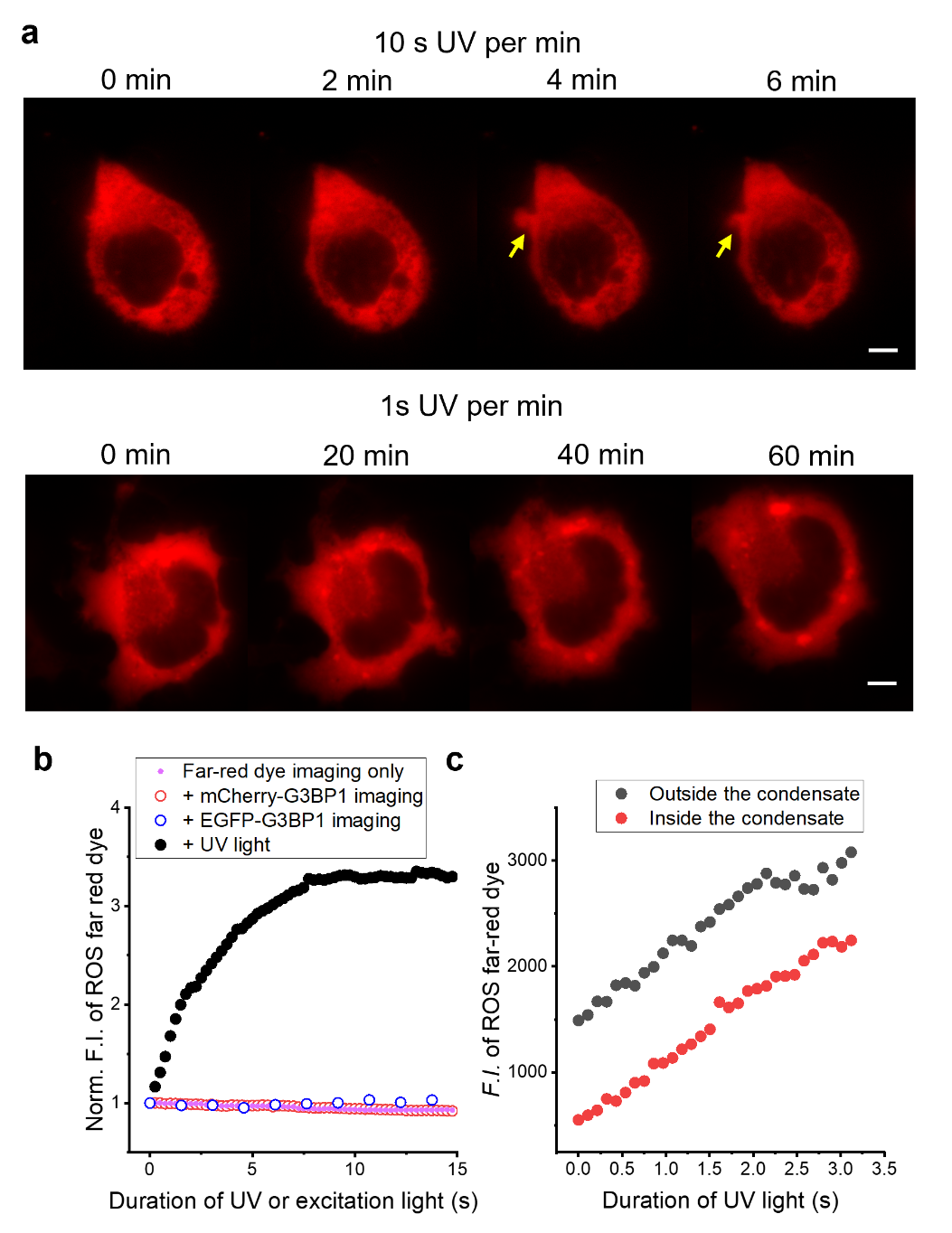

**Figure S7**: Effect of UV irradiation on HEK293T cells transfected with mCherry-G3BP1 and labeled with a far-red ROS dye.

a) Fluorescence images of mCherry-G3BP1. Upper: rapid formation of membrane blebs (yellow arrow) under high dosage of UV irradiation. Lower: slow formation of stress granules under low dosage of UV irradiation. b) Change of intracellular ROS level under various imaging conditions as labeled. c) UV-induced increase in ROS level inside (red) and outside (black) of stress granules. The power density of UV light is 0.245 W/cm^2^. Scale bar: 5 μm.

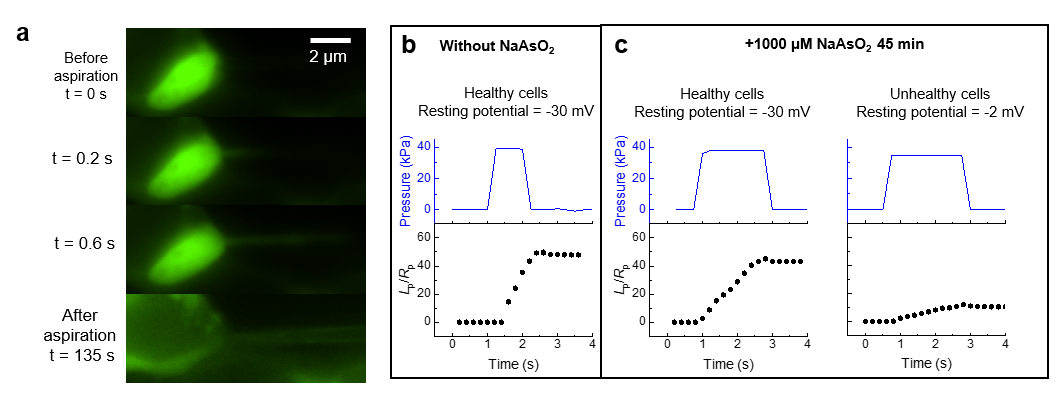

**Figure S8: MAPAC on stress granules under different stress levels**

(**a**) Representative image of the MAPAC of an intracellular stress granule in a HEK293T cell. Frames show the droplet prior to contact, during aspiration, and after; scale bar: 2 µm. (**b**) In untreated cells with a normal resting membrane potential (~-30 mV) the imposed pressure step (blue) produces a rapid deformation (black). (**c**) After NaAsO_2_ treatment, the different behavior among physiologically healthy and unhealthy cells.

**Supplementary Discussion**

The fluorescence (*I*) of a condensate decays exponentially during photobleaching:

$$I=I_{0}e^{-k_{\mathrm{pb}}t} (S1)$$

Here, *I*_0_ is the initial fluorescence, and *k*_pb_ is the bleaching rate.

As observed in Figure 1, the viscosity of the condensate increases exponentially during fluorescence excitation:

$$\eta=\eta_{0}e^{k_{s}t} (S2)$$

Here, *η*_0_ is the initial viscosity of the condensate, and *k*_s_ is the measured solidification rate.

In Figure 2e, we established a linear relation between *k*_s_ and *k*_pb_:

$$k_{s}=S_{L} \phi k_{\mathrm{pb}} (S3)$$

with $\phi$ representing the fraction of labeled protein in the condensate, *S*_L_ represents the slope in Figure 2e.

We note that both equations S2 and S3 are empirical, additional factors such as imaging conditions (e.g., light power, buffer redox potential), the chemistry of the fluorophore, composition and size of the condensate could affect these relations. Notably, photobleaching is not always equivalent to ROS generation, as evidenced by the ability to do FRAP in the presence of ROS scavengers (Figure 3f) and the apparent saturation of *k*_s_ with light power (Figure 1g). Furthermore, the same fluorophore can lead to different rates of solidification in different condensates, as suggested by the effect of unlabeled αSyn on the solidification of Synapsin condensate (Figure 2b).

Combine (S1), (S2), and (S3):

$$\ln\frac{\eta}{\eta_{0}}=k_{s}t= S_{L} \phi k_{\mathrm{pb}}t=S_{L} \phi\ln\frac{I_{0}}{I} (S5)$$

Or,

$$\frac{\eta}{\eta_{0}}=\left( \frac{I_{0}}{I} \right)^{S_{L} \phi} (S6)$$

In Figure 2e, *S*_L_ = 171 ± 12. Therefore,

$$\frac{\eta}{\eta_{0}}=\left( \frac{I_{0}}{I} \right)^{\left( 171\pm12 \right) \phi} (S7)$$

A quick guide to choosing protein label fraction: If the condensate is fully labeled ($\phi$ = 1), 5% bleaching in fluorescence would correspond to a ~4000-fold (1.05^171^) increase in condensate viscosity. If the condensate is 10% labeled ($\phi$ = 0.1), 5% bleaching corresponds to a 2.3-fold (1.05^17.1^) increase in condensate viscosity. If the condensate is only 1% labeled ($\phi$ = 0.01), 5% bleaching corresponds to less than 9% increase (1.05^1.71^) in condensate viscosity.
